# An MRI protocol for anatomical and functional evaluation of the California sea lion brain

**DOI:** 10.1101/2020.08.02.233429

**Authors:** Peter Cook, Vanessa Hoard, Sudipto Dolui, Blaise Frederick, Richard Redfern, Sophie Dennison, Barbie Halaska, Josh Bloom, Kris Kruse-Elliott, Emily Whitmer, Emily Trumbull, Greg Berns, John Detre, Mark D’Esposito, Frances Gulland, Colleen Reichmuth, Shawn Johnson, Cara Field, Ben Inglis

## Abstract

We describe a research MRI protocol for *in vivo* evaluation of pinniped brains using standard human clinical MRI hardware and pulse sequences. Our intended application is to study development of California sea lions (*Zalophus californianus*) exposed *in utero* to domoic acid (DOM) produced by harmful algae blooms in the coastal Pacific Ocean. In cases where the fetus survives to birth, exposure to the toxin *in utero* could result in developmental abnormalities leading to neurological and behavioral deficits. Prior studies on sea lions naturally exposed to DOM as adults have demonstrated hippocampal atrophy and altered mesial temporal connectivity. This MRI protocol is therefore intended to depict the hippocampal formation as the primary region of interest, and to provide longitudinal measures of brain functional and structural connectivity as well as quantitative anatomical evaluations. Scan quality and utility are assessed by comparison with prior studies in live and post mortem sea lion brains. We include the first determination of cerebral blood flow mapping using MRI, and also the first fiber tractography using diffusion-weighted imaging from a live sea lion brain. The protocol also facilitates screening for common neurological pathologies, including tumors, trauma and hemorrhages. We believe the protocol would be suitable for any pinniped that can fit inside a human MRI scanner.

## Introduction

Domoic acid (DOM) is a potent excitatory neurotoxin produced naturally by some species of marine diatoms in the genus *Pseudo-nitzschia*. It accumulates in the viscera of filter-feeding finfish and shellfish such as anchovies, sardines, crabs, clams, mussels and oysters (Bejerano *et al*. 2008). Consumption of fish containing DOM is a major health hazard for marine mammals in particular. The severity and extent of DOM-producing harmful algal blooms (HABs) are increasing worldwide, leading to increased potential for both marine mammal and human exposure to DOM (McCabe *et al*. 2016, Bates *et al*. 2018).

Sea lions have emerged as a focal species for the study of naturally-occuring DOM in the environment, with thousands of California sea lions (CSLs) stranding with neurotoxic symptoms since the 1980s. Research on rehabilitating sea lions has confirmed that high levels of DOM in pregnant mothers can be transferred (with potentially amplified effects) to their pups *in utero* and via lactation, and several case studies indicate that maternal exposure can result in devastating health effects expressed much later in development. We are planning a longitudinal study to identify etiology, symptomatology, and pathology of prenatal exposure to DOM. Here, we describe a detailed MRI protocol for application to sea lions between 6 months and 4 years of age. Our objectives are to provide accurate morphometry of brain structures thought to be at highest risk from DOM toxicosis, to map structural and functional connectivity between the highest risk regions and the rest of the brain, and to define more sensitive and specific clinical criteria for diagnosing DOM toxicosis in wild animals that strand.

### Domoic acid toxicosis

Most neurotoxicity of DOM arises via kainate subtype receptors, although there is some DOM affinity for all classes of ionotropic glutamate receptors (Pulido, 2008; Costa *et al*., 2010). Domoic acid is a kainic acid (KA) analog and has a high binding affinity for KA receptors, triggering a cellular influx of calcium ions and concomitant release of glutamate (Berman *et al*., 2002) which in turn activates N-methyl D-aspartate (NMDA) receptors to cause even greater glutamate release. Rapid influx of intracellular calcium produces efflux of glutathione, an antioxidant, and accumulation of reactive oxygen species in mitochondria. Collectively, these events lead quickly to necrotic cell death (Giordano *et al*., 2006). Across all brain regions, hippocampal and amygdala neurons appear to be at elevated risk due to the density of glutamatergic receptors.

Domoic acid toxicosis was first confirmed in California sea lions in 2000 following a mass stranding event in Monterey Bay in 1998 (Scholin et al., 2000). Acute symptoms included seizures, head weaving, scratching and vomiting. Chronic exposure to DOM produces an epileptic syndrome in sea lions characterized by behavioural changes, seizures and atrophy of the hippocampal formation (Gulland *et al*. 2002, Goldstein *et al*. 2008, Thomas *et al*. 2008, Buckmaster *et al*. 2014). Histology shows hippocampal sclerosis, especially of dentate gyrus and pyramidal cells of CA1, CA3 and CA4, along with gliosis (Scholin *et al*., 2000; Gulland *et al*., 2002; Silvagni *et al*., 2005), and lesions are frequently unilateral (Buckmaster *et al*. 2014).

Domoic acid crosses the intact adult blood-brain barrier slowly, at about the same rate as sucrose, suggesting diffusion-limited transfer rather than an active transport mechanism. However, controlled experimental studies with laboratory mice have demonstrated that DOM can cross the placenta and the mammary gland, causing developmental effects in young rodents (Dakshinamurti *et al*. 1993, Xi *et al*. 1997, Maucher and Ramsdell 2005, 2007). Toxicity models in rodents have been extrapolated to predict similar toxicity from *in utero* exposure in California sea lions (Ramsdell and Gulland 2014), but studies to confirm these effects have not been conducted. Ongoing studies with laboratory primates suggest delayed developmental effects occur in primates exposed *in utero*, but these findings will take several years to confirm, and due to expense, are of limited sample sizes (Burbacher *et al*. 2019, Petroff *et al*. 2019).

Given that DOM readily crosses the placenta (Brodie *et al*. 2006, Costa *et al*. 2010), it is an open question whether the young sea lions that strand with neurological symptoms have been exposed to DOM-contaminated fish while foraging, or if they are suffering from developmental disabilities arising from prenatal exposure. In pregnant sea lions, DOM has been detected in maternal urine, in fetal gastrointestinal contents and urine, and in amniotic fluid, demonstrating that DOM can be transferred to developing fetuses *in utero* (Brodie *et al*. 2006). Recent research indicates that DOM concentrates in maternal amniotic fluid during pregnancy as the fetal fluids act as a “sink” and continually expose the developing sea lion fetus (Lefebvre *et al*. 2018). Domoic acid has also been detected in the milk of sea lion mothers during DOM-producing algal blooms (Rust *et al*. 2014). Furthermore, *Pseudo-nitzschia* blooms typically occur during the spring and summer months off the central and southern California coast, which coincides with the pupping and breeding season of California sea lions. Thus, exposure of pups to DOM may occur *in utero*, during lactation and during early foraging after weaning at nine months.

### MRI of DOM toxicosis

Prior MRI scans at 1.5 tesla (T) revealed considerable abnormal anatomy. Of forty-two sea lions with suspected DOM toxicosis, MRI revealed hippocampal atrophy in 41 animals, accompanied by thinning of the parahippocampal gyrus in 28 animals, increases in ventricle (temporal horn) size in 34 animals, and hyperintensity consistent with gliosis in the parahippocampus in 19 animals (Goldstein *et al*. 2008). The scan protocol included two-dimensional (2D) proton density-weighted, T_2_-weighted and T_2_-weighted FLAIR (fluid-attenuated inversion recovery) scans, plus a three-dimensional (3D) T_1_-weighted gradient echo scan. Lesions were best detected in the T_2_-FLAIR scan. We therefore include a 3D T_2_-FLAIR for clinical evaluations in the present protocol, with the 3D format allowing arbitrary plane selections *post hoc*, while scanning with isotropic spatial resolution in three dimensions is better suited to quantitative morphometry.

Detailed anatomical imaging of the CSL brain was also performed by Montie *et al*. (2009, 2010) to establish healthy baselines of living individuals. They found T_2_-weighted images were preferable to T_1_-weighted scans for identifying normal brain structures. We therefore include a T_2_-weighted scan for structure determination and comparison with a prior atlas (Montie *et al*. 2009).

Recently, Cook *et al*. (2018) used tractography from diffusion MRI (dMRI) to investigate the effects of DOM on hippocampal connectivity in a small sample of postmortem specimens with and without clinical signs of DOM toxicosis. The DOM brains showed clear evidence of reduced hippocampal volume, and potential signs of white matter pathology in the fornix, including reduced fractional anisotropy (FA) and increased mean diffusivity (MD). In addition, these brains showed increased white matter connectivity between the hippocampus and thalamus. These findings mirror those in humans with mesial temporal lobe epilepsy (mTLE) (Concha *et al*., 2010; Dinkelacker *et al*., 2015; Otte et al., 2012), and add support to the idea that wild sea lions with DOM toxicosis might be used as natural models for studying the effects of epilepsy (Buckmaster *et al*., 2014; Ramsdell & Gulland, 2014). We therefore include dMRI in the protocol, and present here a preliminary report of structural connectivity from a live CSL brain.

Behaviorally, disruptions in habituation and sensitization, as well as in spatial memory, have been documented in CSLs with hippocampal atrophy associated with DOM toxicosis. In a sample of 27 wild sea lions undergoing rehabilitation, Cook *et al*. (2016) documented behavioral perseveration in animals with reduced ventral hippocampal volume as measured by MRI. This result was in line with studies showing altered attention and reactivity in rodents with laboratory exposure to DOM (Arkhipov *et al*., 2008; Zuloaga *et al*., 2016) and may be related to observations of repetitive behaviors in DOM-exposed sea lions (Goldstein *et al*., 2008). In another sample of 30 wild sea lions, Cook *et al*. (2015) found altered performance in a T-maze and a daily foraging task related to reduced dorsal right hippocampal volume, which indicated impaired allocentric memory. The same hippocampal region is involved in allocentric spatial memory in humans (Eichenbaum *et al*., 2016). A subset of 11 sea lions from these two studies also underwent functional MRI (fMRI) at 1.5 T. Evidence of reduced functional connectivity between the hippocampus and thalamus was found in animals with DOM toxicosis and resultant hippocampal atrophy (Cook *et al*. 2015). We include a similar fMRI scan in the current protocol, seeking to exploit the greater blood oxygenation level-dependent (BOLD) contrast at 3 T than the prior study at 1.5 T.

### MRI of status epilepticus

Epileptic seizures are common in sea lions that strand with likely exposure to DOM (Goldstein *et al*. 2008). In general, these symptoms are associated with atrophy of the hippocampal formation and are consistent with mTLE. Epileptogenic pathology can be difficult to detect with MRI, and high resolution and contrast are required to delineate the hippocampal formation. Our protocol takes into account many of the recommendations of a consensus report on epilepsy in veterinary medicine (Rusbridge *et al*. 2015) and a suggested protocol for detection of epileptogenic lesions in outpatient human studies (Wellmer *et al*. 2013). In addition, we propose to map the cerebral blood flow (CBF). The CBF is usually considered to be tightly coupled with cerebral glucose metabolism (Roy and Sherrington 1890, Newberg *et al*. 2005). Abnormally high regional CBF (rCBF) has been detected in humans in the peri-ictal period (Nguyen *et al*. 2010, Yoo *et al*. 2017), including in newborns (Mabray *et al*. 2018). In the interictal periods, however, epilepsy is often characterized by reduced CBF in the epileptogenic regions (Detre and Alsop 1999, Wolf *et al*. 2001). Patients with temporal lobe epilepsy were found to have asymmetric rCBF whether or not there was evidence of hippocampal sclerosis, although the patterns of rCBF abnormalities were different with sclerosis (Guo *et al*. 2015).

Arterial spin labeling (ASL) MRI offers a non-invasive measure of rCBF with 2-3 mm spatial resolution, albeit with some caveats. Extant sequences are optimized for the vascular distances, transit delays and geometry of the human brain, and the kinetic models used to compute rCBF rely on assumptions that may not hold in other species. Anesthesia further complicates the picture. To address these issues, we have performed empirical tests to determine whether reasonable CBF mapping in pinnipeds is feasible, and we report preliminary evidence to suggest that sea lion vasculature and blood flow dynamics are sufficiently similar to humans to permit use of human ASL methods. In CSLs with suspected chronic DOM toxicosis, we predict that abnormal rCBF might be detected before significant hippocampal atrophy is observed on anatomical scans, and this should allow us to better define the pathological sequelae.

### Protocol goals and constraints

To summarize, the scan protocol for studying a developing sea lion brain comprises three types of 3D anatomical MRI, functional MRI, a perfusion (CBF) mapping scan, and diffusion MRI. Together, these scanning approaches should facilitate a holistic assessment of the neural effects of DOM toxicity. The entire protocol is constrained by the allowable time under anesthesia, for which we use 1 hour as a guideline because it has been well tolerated by sea lions previously, including animals under a year old. Allowing 15 minutes for intubation, positioning and final health check before the scan commences yields a target maximum protocol duration of 45 minutes, which must include localizer scans and other scanner adjustments.

## Methods

### MRI protocol development

The protocol was developed at the Henry H. Wheeler, Jr. Brain Imaging Center (BIC) at UC Berkeley, on a 3 T Siemens TIM/Trio MRI scanner running version B17A software. Initial tests were performed on post mortem (PM), formalin-fixed sea lion brains provided by The Marine Mammal Center (TMMC), Sausalito CA. A 32-channel receive-only head RF coil was used to establish the spatial parameters required to avoid signal aliasing and approximate scan duration limits commensurate with image signal-to-noise ratios (SNR). The pilot protocol was then applied on PM brains using a 4-channel human neck RF coil in anticipation of scanning live animals. The neck coil (dimensions: L=190 mm, W=330 mm, H=332 mm) is a ring shape, fully open back and front, that is suitable for intubated live animals. The neck coil was previously used in a study of trained dogs (Berns *et al*. 2017).

An issue identified during pilot scanning on PM specimens was the relatively poor performance of acceleration by under-sampling of k-space when using the 4-channel neck coil. The four loops of the array – two loops in the upper half, two in the lower half - offer limited spatial information for reconstruction methods such as GRAPPA (Griswold *et al*. 2002) compared to a 12-channel or 32-channel array coil as for human brain imaging. Accordingly, we made the decision to use full k-space sampling without acceleration in most live animal scans. An exception was the T_2_-weighted 3D anatomical scan, which would have required a prohibitively long scan duration without acceleration. We also opted not to use simultaneous multi-slice (SMS) acquisition for fMRI or dMRI, again because of the risk of artifacts. Acceleration methods might be included at a later date, if a larger phased-array coil is available to replace the 4-channel neck coil.

### Animal procedures

Scanning was performed on five California sea lions (CSL) undergoing rehabilitation at TMMC. All animals either had symptoms consistent with DOM toxicosis or other neurological conditions. Scans were also performed on two CSLs housed at Six Flags Discovery Kingdom (SFDK) in Vallejo, CA. The SFDK animals comprised one 3 year-old male adopted from TMMC after repeated strandings that was subsequently diagnosed with epilepsy, and one captive-bred 7 month-old pup. The TMMC animals were treated under an authorization granted to the Marine Mammal Health and Stranding Response Programme by the National Marine Fisheries Service scientific research permit 18786-04, and approved by the Animal Care and Use Committee (ACUC) at TMMC. Procedures on the SFDK animals were approved by the ACUC of SFDK. The Office of Laboratory Animal Care (OLAC) at UC Berkeley determined that no formal oversight was required by the Berkeley ACUC since the scans were clinical evaluations performed as part of the animals’ normal veterinary care under the authorization of TMMC or SFDK, and with all medical procedures and welfare monitoring conducted by their veterinary staff. Key data on all animals is included in Table 1.

**Table 1:**
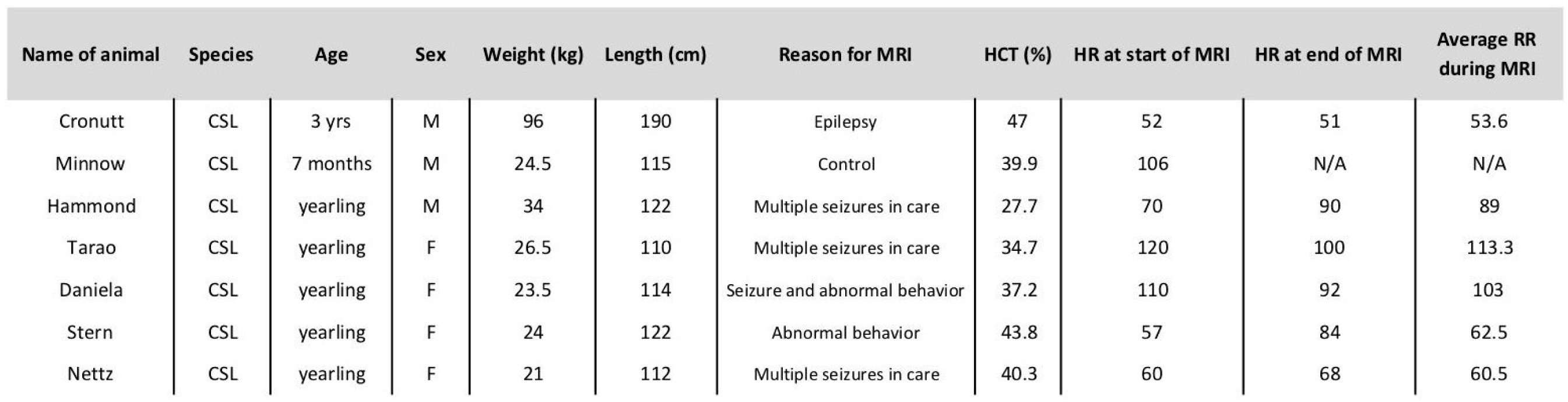
Key data on animal subjects.

| Name of animal | Species | Age | Sex | Weight (kg) | Length (cm) | Reason for MRI | HCT (%) | HR at start of MRI | HR at end of MRI | Average RR during MRI |
| --- | --- | --- | --- | --- | --- | --- | --- | --- | --- | --- |
| Cronutt | CSL | 3 yrs | M | 96 | 190 | Epilepsy | 47 | 52 | 51 | 53.6 |
| Minnow | CSL | 7 months | M | 24.5 | 115 | Control | 39.9 | 106 | N/A | N/A |
| Hammond | CSL | yearling | M | 34 | 122 | Multiple seizures in care | 27.7 | 70 | 90 | 89 |
| Tarao | CSL | yearling | F | 26.5 | 110 | Multiple seizures in care | 34.7 | 120 | 100 | 113.3 |
| Daniela | CSL | yearling | F | 23.5 | 114 | Seizure and abnormal behavior | 37.2 | 110 | 92 | 103 |
| Stern | CSL | yearling | F | 24 | 122 | Abnormal behavior | 43.8 | 57 | 84 | 62.5 |
| Nettz | CSL | yearling | F | 21 | 112 | Multiple seizures in care | 40.3 | 60 | 68 | 60.5 |

The MRI suite was prepared for CSL scanning by removing extraneous furniture and impediments to the large number of staff required to transport and monitor an animal under anesthesia. Plastic sheeting was laid on all carpeted floor areas for hygiene. All participants received an MRI safety briefing, and plans and procedures were reviewed by senior personnel at the start of each scan session. During a session an experienced BIC staff member was assigned as the duty safety officer and was given the sole task of observing activities to identify the development of any potentially hazardous situations. Access to the magnet room – designated Zone IV according to the American College of Radiology (ACR) safety zones principle - was restricted to designated veterinary care staff, all of whom were fully screened for MRI contraindications. Equipment and supplies required for maintenance of anesthesia and animal monitoring were pre-approved for Zone IV use by BIC staff. All other items were prohibited from Zone IV without the express consent of the duty safety officer.

The CSL transport crate was located outside the scanner suite in a corridor (ACR Zone II) and the anesthesia rig stationed nearby. The lack of an MRI-compatible anesthesia machine and ventilator necessitated induction with the vaporizer in the corridor, followed by relocation to the operator room (ACR Zone III) for the duration of the MRI scan. Gas delivery to the magnet in Zone IV was through a ten meter rubber hose carrying oxygen and inhalant. The hose was run via a waveguide between Zones III and IV. The distance from waveguide to magnet center is comparable to the distance from the waveguide to the corridor, allowing the use of the same hose for anesthesia induction and maintenance with minimal delay for switching.

Animal handling and anesthesia procedures were established by veterinary staff of TMMC and SFDK in consultation with a veterinary anesthesiologist (KK-E). Animals were fasted overnight to ensure an empty stomach. If necessary, an animal was sedated via intramuscular injection of midazolam (0.1-0.3 mg/kg), butorphanol (0.1-0.3 mg/kg) and medetomidine (0.1-0.3 mg/kg) via syringe while the animal was still in the transport crate. Once the animal was sedated, the crate door was opened and a conical face mask fitted over the snout. Anesthesia was induced with 5% isoflurane in oxygen. As soon as jaw tone was absent, the animal was removed from the crate onto a towel or tarpaulin on the floor, then lifted onto the foam pad of an MRI compatible gurney. The foam pad was then lifted onto the gurney at its lowest height for endotracheal intubation (Figure 1). Following intubation, animals were maintained at an appropriate level of isoflurane and, at the lead veterinarian’s discretion, the sedative medetomidine was reversed with atipamezole. The MRI compatible gurney was found to be useful for safe and efficient animal handling. The initial lift of approximately 30 cm onto the foam pad was easily and safely achieved using a tarpaulin for the largest animal scanned so far (96 kg). The gurney’s height could then be lifted to its maximum extent with the animal completely supported, obviating the need to lift or rotate the animal unsupported for the remainder of the session.

**Figure 1.**
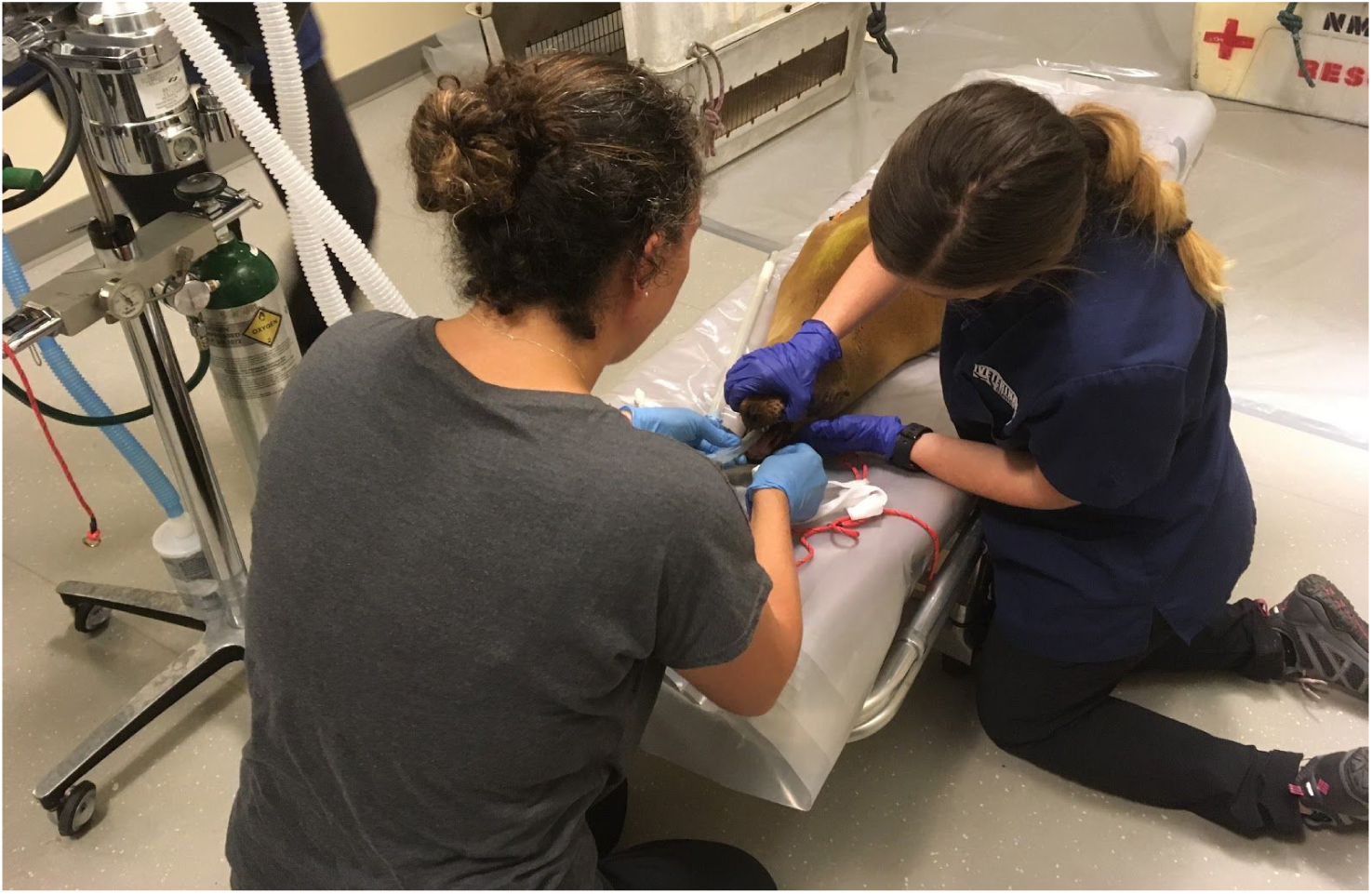
A sea lion being intubated on an MRI compatible gurney located in Zone II, a corridor outside the MRI scan suite.

Animals were scanned in ventral or dorsal recumbency (equivalent to human prone or supine positioning, respectively), depending on size and additional procedures such as blood draw from vena cava. Dorsal recumbency was preferred because it allowed the easiest vital signs monitoring. The recumbency was found to have no significant effect on MRI scanning. For transit from the corridor into the scanner room it was necessary to temporarily disconnect the anesthesia tubing from the endotracheal tube, reconnecting once the gurney was wheeled into Zone IV and the vaporizer wheeled from the corridor into Zone III. The MRI patient bed height was set to match the gurney height so that the pad supporting the animal could be transferred to the patient bed directly, on top of the standard MRI bed padding. This placed the animal’s head at an appropriate height for the receiver coil (Figure 2). Once the CSL was on the MRI patient bed, it was only necessary to slide the animal along the pad to center the brain in the receiver coil; this could be performed conveniently and safely at the magnet bore. Two nylon straps affixed to the MRI bed were used to restrain the animal in case of seizures or unintended recovery from anesthesia while in the magnet. A pediatric fiberoptic pulse-oximetry sensor (Nonin 8600) was placed on the animal’s tongue to measure heart rate (HR) and arterial blood oxygen saturation. However, intermittent signal meant that vital signs were primarily determined by direct observation by a technician permanently stationed in Zone IV. Foam pads were placed either side of the animal’s head, to provide hearing protection as well as restrain the head in the appropriate position in the receiver coil. We opted not to use ear plugs because of the potential difficulty removing them in emergency situations. Anesthesia was maintained with ∼2% isoflurane in oxygen, the isoflurane level adjusted based on vital signs. Animals were intubated but breathed spontaneously. A plastic disposable pediatric circle system with CO_2_ absorber used as a semi-closed circle system and activated charcoal isoflurane scavenger (F/Air, AM Bickford, Wales Center, NY, USA) were mounted on an MRI compatible stand next to the magnet bore (Figure 3). The circle system and scavenger canister are plastic and determined to be MRI compatible by BIC staff. Monitoring included visual counts of breathing rate and measurement of HR using a disposable plastic stethoscope (Proscope 665, American Diagnostic Corporation, Hauppauge NY, USA). Animals were cooled using the magnet bore fan and plastic bags of ice placed on the rear flippers. Lack of an MRI compatible thermometer meant that rectal temperature could only be taken outside Zone IV, before and after scanning. Hypercapnia was avoided with intermittent manual ventilation using a 3L rebreathing bag. Past experience has shown that manual ventilation at a rate of 1/minute is sufficient to aid elimination of CO_2_ in anesthetized sea lions.

**Figure 2.**
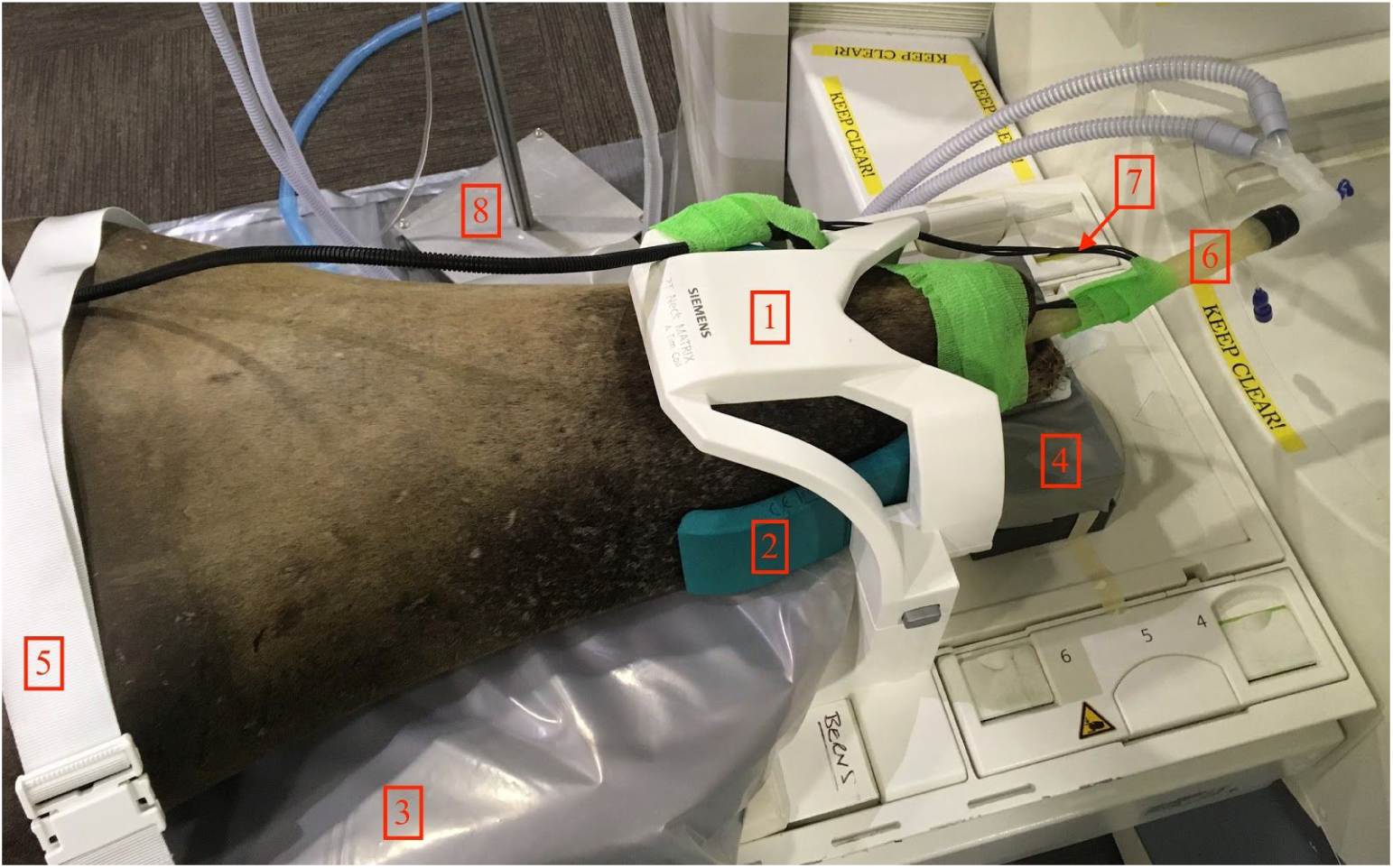
A 3 year-old California sea lion positioned on the patient table for scanning. 1: Receive-only neck coil. 2: Foam padding for head restraint and hearing protection. 3: Pad from the MRI compatible gurney, wrapped in plastic sheeting for hygiene, and used to lift/support the animal throughout. 4: Foam to support/position the snout. 5: Nylon strap for restraint. 6: Endotracheal tube and anesthesia line. 7: Pulse oximeter fiber optic cable. 8: The base of the non-magnetic stand used to hold the anesthesia circle system.

**Figure 3.**
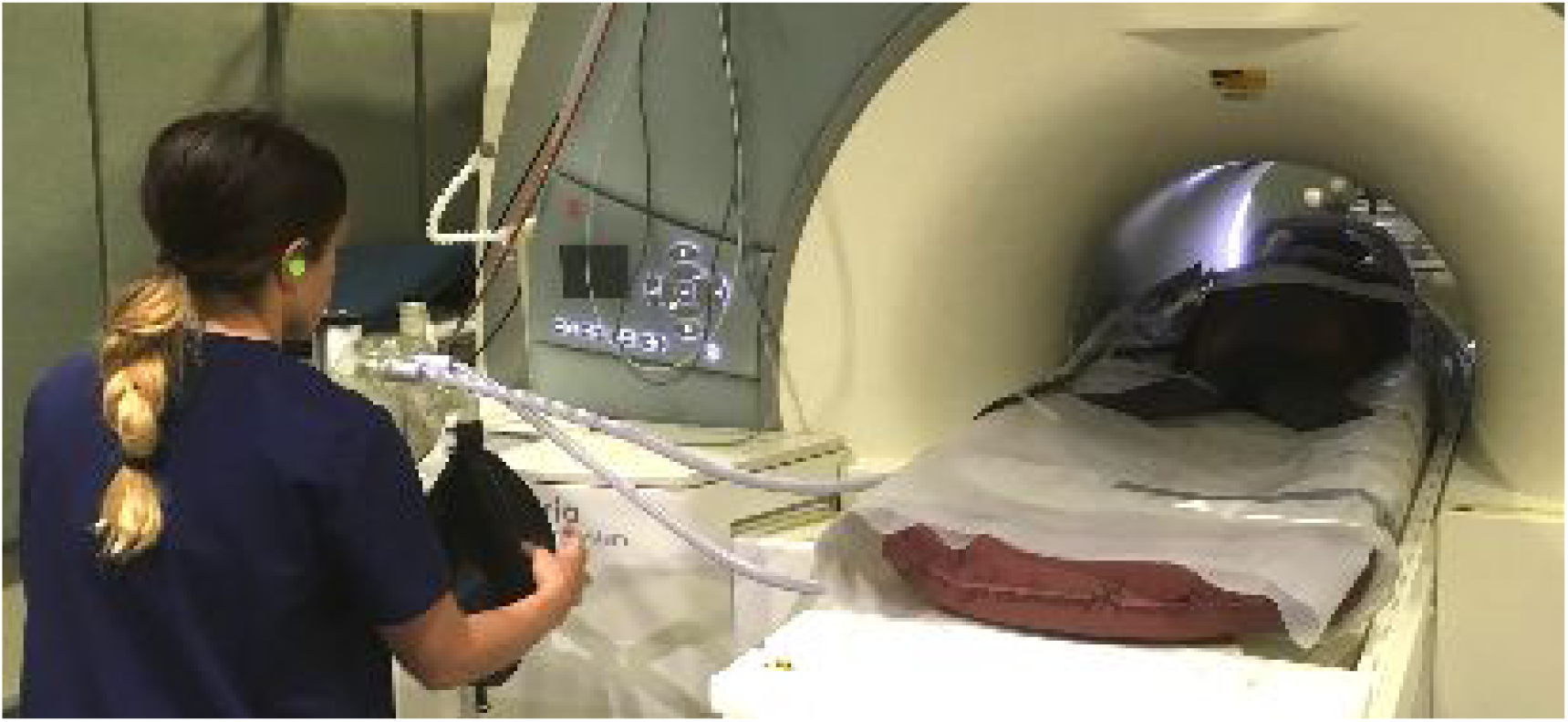
A pediatric circle system with CO_2_ absorber mounted at the magnet bore, with the sea lion supported inside the magnet on the gurney pad. An isoflurane scavenger is connected below the anesthesia circle system (not visible). The veterinary technician remains in Zone IV to monitor vital signs directly and periodically assist ventilation manually via a 3L rebreathing bag.

At the completion of MR scanning, the gas line was temporarily disconnected from the endotracheal tube and the animal transferred via the gurney to the transport crate in Zone II. Once in the crate, 100% oxygen was supplied until the animal showed signs of alertness, when the endotracheal tube was withdrawn and the door of the crate closed. Respiratory rate and character were monitored during recovery. If not already reversed, medetomidine was reversed with atipamezole. Ice was added to the crate to maintain body temperature.

### T_1_-weighted 3D MP-RAGE

Initial testing with a sagittal slab-selective 3D magnetization-prepared rapid acquisition of gradient echo (MP-RAGE) sequence, a conventional scan for human brain research, exhibited pronounced respiratory artifacts. The soft tissues of a sea lion’s neck and lower jaw, which are both within the 3D field-of-view, move slightly with the chest during respiration. We opted to change the prescription from sagittal to dorsal. This has the effect of swapping the phase encoding axes in the 3D acquisition. In the dorsal prescription, the first phase encoding direction is then left-to-right (L-R) and the second phase encoding direction is anterior-to-posterior (A-P) (equivalently ventral-dorsal). The slower k-space sampling A-P is more robust to throat movement at the respiratory rate. Dorsal 3D imaging with slab-selective excitation retains head-to-foot (H-F) frequency encoding, thereby permitting digital filtering of extraneous signal from the neck and upper body to eliminate aliasing. Remaining acquisition parameters for the MP-RAGE were adopted from the third phase of the Alzheimer’s Disease Neuroimaging Initiative (ADNI-3) (Weiner et al. 2017). We have found the combination of repetition time, inversion time and flip angle used in ADNI-3 gives excellent gray/white tissue contrast in human brains. In accordance with standard clinical and research practice, we used the Prescan Normalize option to correct for the bias field imposed by the neck receiver coil. We opted to acquire standard 1 mm isotropic voxels, although we also tested 0.8 mm x 1 mm x 1 mm in two subjects. The higher resolution in the frequency encoding dimension is limited by gradient speed, while the 1 mm x1 mm dimensions of the two phase-encoded dimensions were limited by scan time constraints. Acceleration using GRAPPA produced unacceptable banding artifacts with the 4-channel neck coil. On balance, the benefits of pushing beyond (1 mm)^3^ were marginal for a scan of approximately 7 minutes duration.

### T_2_-weighted 3D SPACE

Previous scanning of live sea lions has generally been conducted using 2D multiple spin echo sequences with two separate TEs per scan: a short TE of around 14 ms for a minimally T_2_-weighted scan (commonly referred to as proton density (PD) contrast) for visualization of tissues, and a longer TE of around 100 ms for visualization of fluid-filled cavities. These scans have diagnosed the presence of gas-filled bubbles (Van Bonn et al. 2011, 2013) and have been used for volumetric measurements of live CSL brain (Montie et al. 2009, 2010). An issue for quantification of small structures is the relatively thick slices used in 2D imaging, however. Voxel dimensions of 0.3 mm x 0.3 mm x 2.5 mm are typical. For research purposes, higher resolution through plane can be useful for tissue segmentation, e.g. for hippocampal volumes. In combination with a high-resolution MP-RAGE, a high resolution T_2_-weighted 3D scan may permit tissue typing and quantitative morphometry. We thus decided that 3D imaging was more likely to be useful in a longitudinal study of DOM toxicosis, and we focused on a long TE to compliment the structural information obtained in the MP-RAGE.

There are several approaches to multiple spin echo imaging, as reviewed by Mugler (2014). Variants include fast spin echo (FSE), turbo spin echo (TSE) and rapid acquisition with relaxation enhancement (RARE). Multiple lines of k-space are acquired per excitation, but in general only one line of k-space is acquired per refocusing period. The effective TE, which by convention arises at the center of k-space, can be set to short or long values as desired. The number of refocusing periods (spin echoes) per excitation, the flip angle used for each refocusing pulse, and the effective TE at the k-space center all have profound effects on both image contrast and artifact levels. Imperfections in the refocusing radiofrequency (RF) pulses lead to complex pathways for magnetization, making artifacts likely. This has led to the development of variable flip angle (VFA) schemes across a single set of spin echo acquisitions, plus tactics to reduce artifacts that remain. On the Siemens platform, the SPACE sequence (Sampling Perfection with Application optimized Contrasts by using different flip angle Evolutions) has been optimized for 3D T_2_-weighted imaging of the brain. A remaining problem using VFA refocusing is produced by structures which have short T_1_ compared brain tissue, such as subcutaneous lipids. Short T_1_ regions may appear mottled. The artifact, which arises from non-refocused signal, can be minimized using the principle of phase cycling of two otherwise identical scans. Alternatively, the relative contribution of non-refocused signal to properly refocused signal can be reduced by shortening the spin echo train, e.g. breaking it into two separate acquisitions. Either tactic doubles the overall acquisition time, however. We found that the phase-cycling approach with two averages at TE=408 ms gave better image quality and preferable T_2_-weighted contrast than splitting the echo train into two halves, which limited the TE to a maximum of 256 ms.

In common with the MP-RAGE, we used a sea lion dorsal prescription with the first phase encoding axis set L-R to minimize respiratory ghosts. Given the historical use of 2D images with high in-plane resolution to depict the hippocampal formation in sea lions, we decided to acquire the smallest isotropic voxels possible. Voxel dimensions of (0.8 mm)^3^ retained excellent contrast-to-noise ratio (CNR) and minimal artifacts. However, an issue with any T_2_-weighted 3D sequence is the long scan duration. We used GRAPPA with an acceleration factor 2 in the second dimension (first phase encode axis), and 5/8ths partial Fourier in the third dimension (second phase ecode axis) to bring the acquisition time down to 6 minutes. Banding from GRAPPA, even with the relatively poor performance of a 4-channel neck coil, is less of a problem than artifacts due to VFA refocusing and from short T_1_ species, and a total scan time of 12 minutes would have been prohibitive.

### T_2_-weighted 3D FLAIR

The T_2_-weighted fluid-attenuated inversion recovery (FLAIR) scan is a mainstay of radiological imaging and is preferred for identifying edema and hippocampal atrophy, as may arise with DOM toxicosis. A standard clinical exam uses multi-slice 2D scans with the slice prescription established to cover regions of greatest interest. The inversion recovery (IR) delay, applied once per slice, limits the total number of planes that can be acquired in a reasonable amount of time. While the in-plane spatial resolution can be as high as 0.3 mm, the slice thickness is generally rather coarse and 3-5 mm thick slices are common. For a general examination, and for possible quantitative morphometric uses, a 3D scan permits *post hoc* slicing across any brain structure of interest, albeit at a lower resolution in two of three dimensions compared to the typical 2D scans.

We acquired a 3D T_2_-FLAIR using a sea lion dorsal prescription to avoid respiratory ghosts, as above. The non-selective IR delay for CSF nulling and other parameters were set as for a human brain on the basis that we would expect the T_1_ of CSF to be similar across mammalian species. Unacceptable banding artifacts with GRAPPA acceleration necessitated the use of 6/8ths partial k-space in the 3rd dimension to reduce the scan time, and this produces some Gibbs ringing artifact and partial smoothing in the third (A-P) dimension. As for the MP-RAGE, we utilized the Prescan Normalization to correct the receive bias field.

### Functional MRI

Cook *et al*. (2015) used fMRI under anesthesia - typically referred to as resting-state fMRI because of the absence of an explicit task - to demonstrate altered hippocampal-thalamic connectivity in sea lions with hippocampal lesions likely caused by exposure to DOM. The prior study was conducted at 1.5 T and used an axial slice prescription. Given that the level of image distortion inherent to the echo planar imaging (EPI) sequence is expected to double at 3 T compared to 1.5 T, we opted to test all three cardinal slice prescriptions for BOLD contrast fMRI at 3 T. The voxel dimensions were fixed at (2.5 mm)^3^, which is approaching the maximum performance of the scanner hardware. For CSL Cronutt, the largest animal scanned so far, a dorsal prescription required only twenty-six 2.5 mm slices to cover the entire brain, whereas 30 and 35 slices were required to cover the brain in axial and sagittal prescriptions, respectively. The dorsal prescription thus offers the best coverage, *i.e.* shortest minimum TR.

We next considered the distortion and Nyquist ghosting inherent to the phase-encoded axis of the EPI pulse sequence. While H-F phase encoding produces symmetric distortions dorsally, L-R phase encoding allows faster gradient switching without violating the (human) peripheral nerve stimulation limits that are built-in to the scanner monitoring. (These limits cannot be disabled in a scanner with FDA approval.) Furthermore, for a 190 mm x 190 mm image the H-F phase encoding caused aliasing of signal from the neck to overlap the brain. It also produced Nyquist ghosts from both neck and snout aliased onto the brain (Figure 4, left panel). By comparison, L-R phase encoding produced asymmetrically distorted brain images but there was no signal aliasing and the Nyquist ghosts did not overlap the brain (Figure 4, right panel).

**Figure 4.**
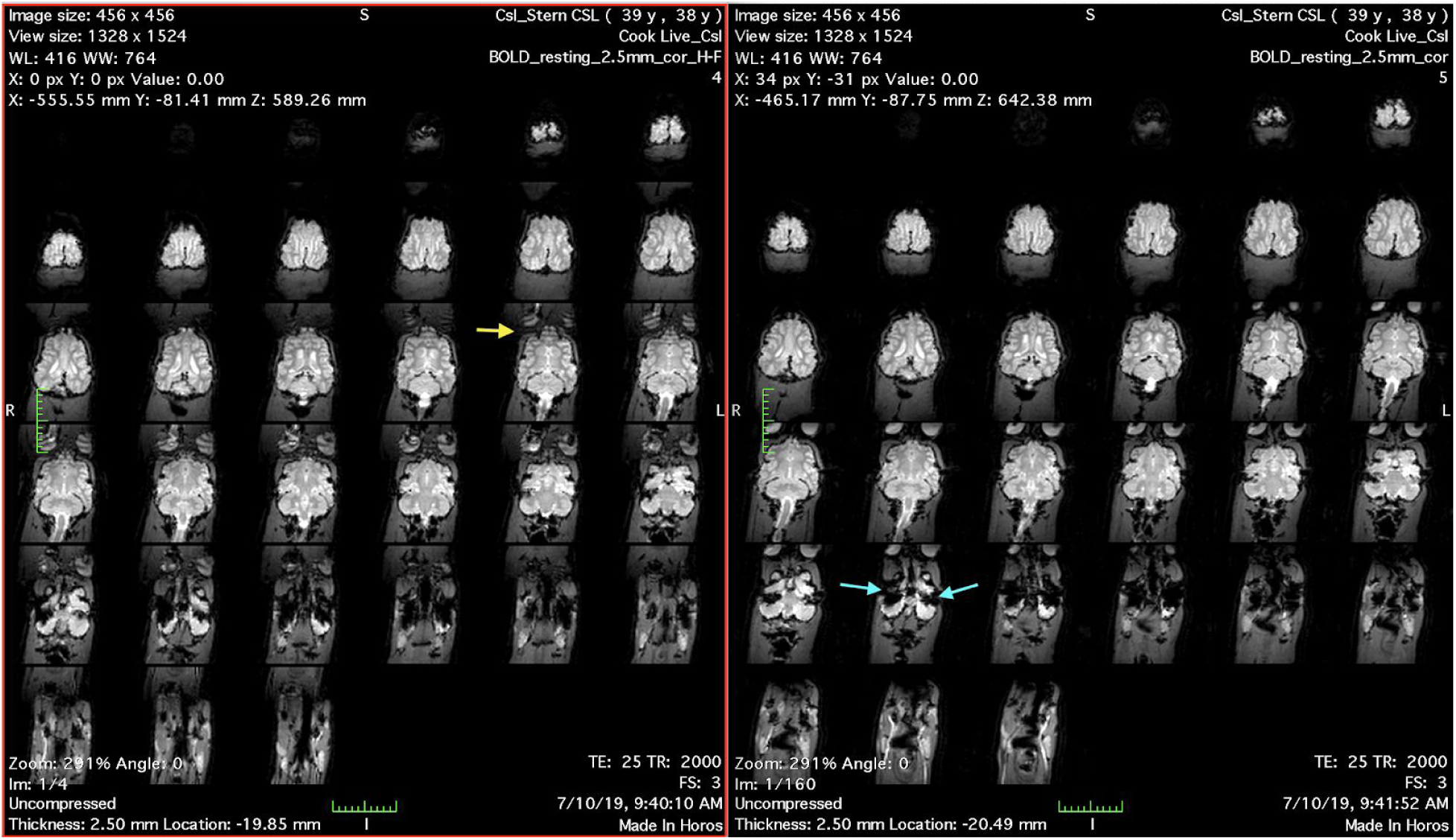
Left: Dorsal sections of BOLD contrast EPI with phase encoding set H-F. Note the banding caused by Nyquist ghosts and aliased signals overlapping the eyes and frontal lobe (yellow arrow). Right: Dorsal EPI with phase encoding set L-R, eliminating the banding artifact. Regions of dropout in cerebellum (blue arrows) were present in all slice prescriptions and regardless of phase encoding direction.

For BOLD contrast we selected TE=25 ms, which is comparable to a TE used for human frontal or temporal lobe fMRI and approximates the T_2_* of brain tissues at 3 T. We did not attempt to quantify regional signal dropout but we noticed bilateral signal losses in cerebellum (Figure 4, blue arrows) regardless of slice prescription, most likely because of the low position of the auditory bullae.

Selecting a suitable duration for task-free fMRI experiments requires some understanding of physiological noise sources, especially changes in arterial CO_2_ which for the human brain tend to fluctuate near to the frequency range most often studied in functional connectivity analyses (Tong & Frederick 2010, Tong & Frederick 2014, Tong *et al*. 2013, Tong *et al*. 2019). We can record chest motion and HR, but given the intermittent reliability of pulse oximetry from the tongue and the limitations of respiratory models that derive arterial CO_2_ changes from chest motion alone, we decided that we would instead rely upon data-driven de-noising methods. A five-minute scan - 160 volumes at TR=2000 ms - should provide adequate compromise between a sensitivity to functional brain networks and ability to characterize and remove physiological noise. We considered testing SMS slice acceleration to decrease TR but were concerned about artifacts which were visible in phantom tests with acceleration factors as low as two. On balance, we decided that the small sampling benefit from halving the TR was not worth the risk of high artifacts in the data. We will revisit the possibilities afforded by SMS acceleration at a later date, when a better RF coil might be available in place of the human neck coil.

### Cerebral blood flow mapping

Arterial spin labeling (ASL) methods are well developed for human brain imaging, but to date they have not been applied to a pinniped. Several ASL variants are available. For humans, the consensus view is that pseudo-continuous ASL (PCASL) is preferred, along with background suppression of static signal (Alsop *et al*. 2015). We have considerable experience with a background-suppressed (BS) stack-of-spirals PCASL sequence (Vidorreta *et al*., 2014, Vidorreta *et al*., 2017) that uses unbalanced gradients during labeling to improve robustness to off-resonance effects and reduce labeling inefficiency due to pulsatility in the carotid arteries (Wu *et al*. 2007, Zhao *et al*. 2017).

Factors that can affect ASL sensitivity and CBF quantification include: (i) the (vascular) distance from the site of arterial blood labeling to the brain tissues, which determines optimal sequence timing, (ii) the blood velocity and pulsatility in the carotid arteries, which affects labeling efficiency, and (iii) the hematocrit, which affects the T_1_ of the labeled blood as used in CBF quantification from raw ASL images. In humans, a labeling period of 1800 ms provides a suitable compromise between sensitivity and efficiency (Alsop *et al*. 2015). Timing of both the labeling period and the post-labeling delay (PLD) are important for accurate quantification of CBF. The time for blood to flow from the labeling position in the neck to the brain tissue is parameterized by the arterial transit time (ATT). The PLD should be sufficiently long that all inverted blood water is delivered to the brain tissue prior to image acquisition. If the PLD is shorter than the slowest ATT then the CBF map can be biased by incomplete delivery of label. In the extreme, bright spots can appear in the final images, representing labeled blood still resident in arteries. On the other hand, the label decays with the T_1_ of blood. Making the PLD too long will degrade the magnitude of the label unnecessarily, leading to low SNR in the perfusion-weighted images.

To adapt the stack-of-spirals PCASL sequence to sea lion brains, we began by mapping the arteries using a time-of-flight (TOF) angiography sequence applied in the sagittal (26 sec scan) and dorsal (32 sec scan) directions, and displaying the maximum intensity projection (MIP) images (Figure 5). While the left and right external carotids are quite far apart and follow the curvature of the neck, there is a location at which we could place the ASL labeling plane approximately perpendicular to the vessels. The labeling plane (Figure 5, hatched bar) was located 90-105 mm caudal to the center of the 3D imaging volume (Figure 5, yellow box), with the imaging volume centered over the brain. The longest distance of 105 mm was used on the largest sea lion scanned so far, 3 year-old CSL Cronutt, and is comparable to the distance used on an adult human brain, consistent with Cronutt being approximately the same size as a large adult human. We used approximately 90 mm for the smaller yearling sea lions and note that in empirical tests on humans we have found that CBF values are reproducible even if the labeling plane location is varied by a few millimeters.

**Figure 5.**
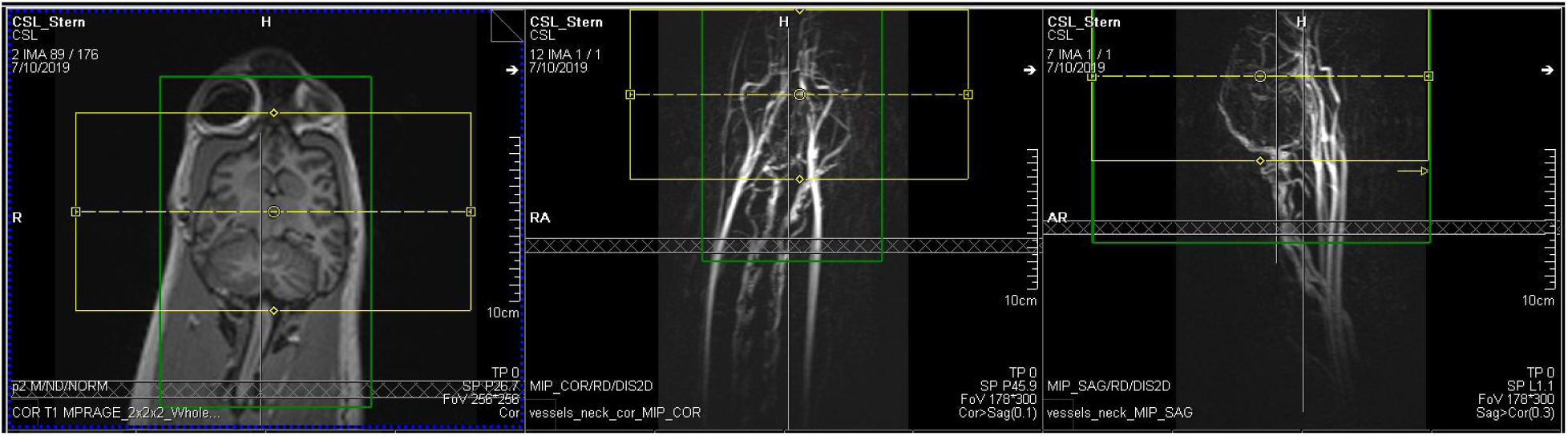
Prescription of spatial parameters for ASL scan on yearling CSL Stern, using a dorsal section through an MP-RAGE to determine the brain extent (left panel) and maximum intensity projections of dorsal (center) and sagittal (right) time-of-flight angiograms to allow correct placement of the labeling plane relative to the carotid arteries. The 3D imaging volume is depicted by the yellow box and the adjustment volume for magnetic field optimization is depicted as a green box. The labeling plane is shown as a horizontal cross-hatched rectangle located 90 mm below the centroid of the imaging volume.

A labeling duration of 1800 ms and PLD of 1800 ms are preferred for the stack-of-spirals PCASL on a normal human brain. A longer PLD of 2000 ms is suggested when vasculature may be compromised, *e.g.* in stroke patients (Alsop *et al*. 2015). However, anesthetized sea lions tend to have a higher HR than awake humans - rates of 80-100 bpm are typical in sea lions - raising the possibility that arterial blood velocity and ATT might be appreciably faster in sea lions than humans and that shorter labeling duration and/or PLD might be warranted. We were also concerned that the hematocrit would be significantly higher than for typical humans, leading to shorter blood T_1_. (Hematocrit values were determined after each scan, from blood drawn at the end of the session, but a value was not available for an individual animal while setting up the PCASL parameters.) We therefore conducted empirical tests of the labeling duration (1200, 1500 and 1800 ms) and PLD (1200, 1500 ms) in different combinations across different animals, as scan time permitted. Given that there is no prior knowledge on which to assess regional CBF in sea lions, we determined that the combination of 1800 ms labeling duration and 1200-1500 ms PLD gave the most reasonable CBF maps, *i.e.* fewest recognizable errors and greatest SNR.

### Diffusion imaging

Abnormal white matter connectivity is expected in animals with hippocampal atrophy caused by DOM toxicosis. Using a diffusion imaging sequence developed specifically for scanning fixed, post mortem specimens, Cook *et al*. (2018) observed signs of white matter pathology in the fornix as well as increased connectivity between the hippocampus and thalamus in sea lions with suspected DOM exposure, compared to brains from animals with no neurological signs of DOM toxicosis. Such a pattern has been observed previously in human patients with mTLE (Concha *et al*., 2010; Dinkelacker *et al*., 2015; Otte *et al*., 2012). Diffusion MRI has also been used to study chronic, low-level DOM exposure in live macaques (Petroff et al. 2019), and preliminary evidence indicates altered white matter parameters in the fornix as well as the internal capsule.

The prior tractography study by Cook *et al*. (2018) was limited by hardware constraints to 1 mm isotropic voxels. Each brain was scanned at 3 T for approximately 8 hours using a novel dMRI sequence that optimizes SNR for post-mortem tissues with short T_2_ (Miller *et al.,* 2012, Foxley *et al*., 2014, Berns *et al*., 2015). For dMRI of anesthetized sea lions, however, total scan time is a more stringent constraint than either hardware limits or even image SNR. Our preliminary tests on PM specimens suggested that (2 mm)^3^ resolution would be possible in an acceptable scan time using b=1000 s/mm^2^ at TE=93 ms and TR=5800 ms. A partial Fourier scheme, 6/8ths of the phase encode direction, was used to reduce TE below 100 ms and maintain adequate SNR. Although higher spatial resolution and/or larger diffusion weighting were supported by the intrinsic SNR of the neck receiver coil, either option would be prohibitively slow for a complete scan protocol of ∼45 minutes. In preliminary tests we also assessed GRAPPA with acceleration factor two in the phase encoding direction, to reduce the TE and halve EPI phase encode distortions, but we observed strong residual aliasing artifacts that might lead to false tracts. We considered SMS acceleration factor two but decided that, this being the first study of live sea lions with dMRI, we should first establish a performance baseline, again in case of residual aliasing (often termed “leakage” for SMS) presenting as false tracts.

The first task in a live animal was to determine optimal slice coverage and evaluate distortions and signal dropout, taking into account our earlier findings from the fMRI scan which also uses an EPI readout. We tested dorsal and sagittal prescriptions with the phase encoding direction set L-R and A-P, respectively. On 3 year-old CSL Cronutt, forty 2-mm slices were sufficient to cover the entire brain dorsally whereas forty-eight 2-mm slices barely covered the brain sagittally. Neither prescription exhibited large regions of static signal loss, and spatial distortions were deemed not to be significantly different by inspection. In both cases, fat presaturation was highly effective at suppressing subcutaneous lipid and blubber signals so that Nyquist ghosts were weak and aliased outside the brain. However, the more efficient slice coverage offered by the dorsal prescription permitted a lower TR and more diffusion-encoded directions per unit acquisition time.

The first full dMRI data set was obtained on CSL Cronutt: thirty diffusion-weighted (DW) directions at b=1000 s/mm^2^ plus three b=0 images, one acquired before each block of ten DW images, to permit improved eddy current correction. The total acquisition time was 3 min 17 sec. Raw image quality on initial inspection appeared to be quite good (Figure 6). Subsequently, on yearling CSL Stern we increased the number of DW directions to 64 plus six b=0 images, retaining b=1000 s/mm^2^, for a total acquisition time of 6 min 52 sec. Finally, on yearling CSL Tarao we attempted to change the acquisition strategy to a two-shell dMRI model (Wu & Alexander, 2007) using fifty DW images at b=500 s/mm^2^ interleaved with fifty DW images at b=1000 s/mm^2^, plus 11 images at b=0, in an acquisition time of 10 min 50 sec, which is around the maximum duration permissible in the scan protocol for an anesthesia period not to exceed an hour. Inspection of the dMRI scans on Tarao revealed a previously overlooked problem. We noticed non-stationary signal voids in some but not all DW directions and in some but not all slices (Figure 7). Assuming there was a problem with the particular directions used in the two-shell scheme, we acquired the 64 DW directions as used for CSL Stern. Yet the non-stationary signal voids were still apparent, again in slice positions that were highly variable.

**Figure 6.**
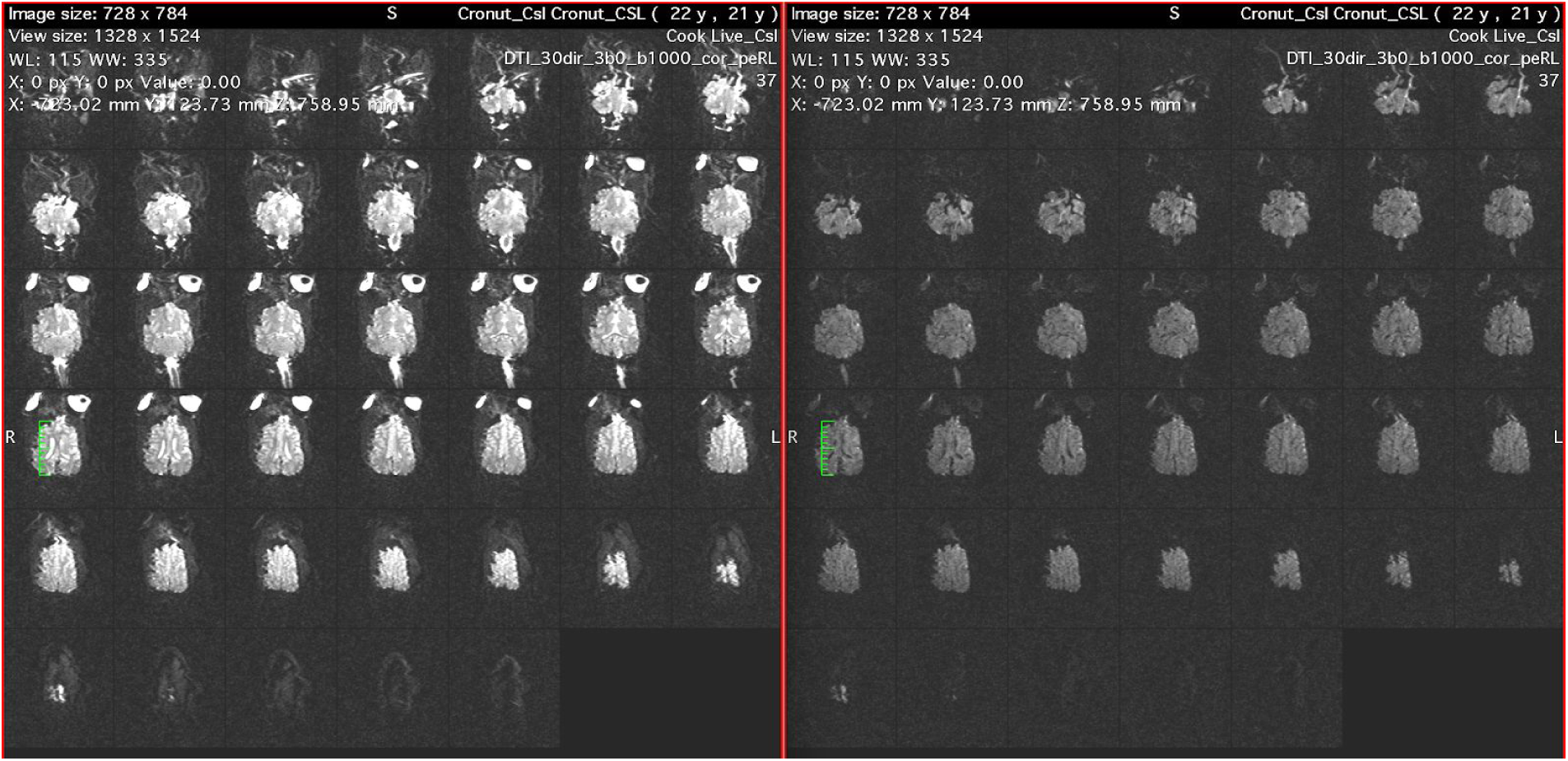
Example diffusion-weighted images acquired from 3 year-old CSL Cronutt at b=0 (left) and b=1000 s/mm^2^ (right). Spatial resolution is (2 mm)^3^. For these dorsal slices the phase encoding is L-R. Note the distortion in the phase encoding direction, most easily recognized in the non-circular signal of the eyes.

**Figure 7.**
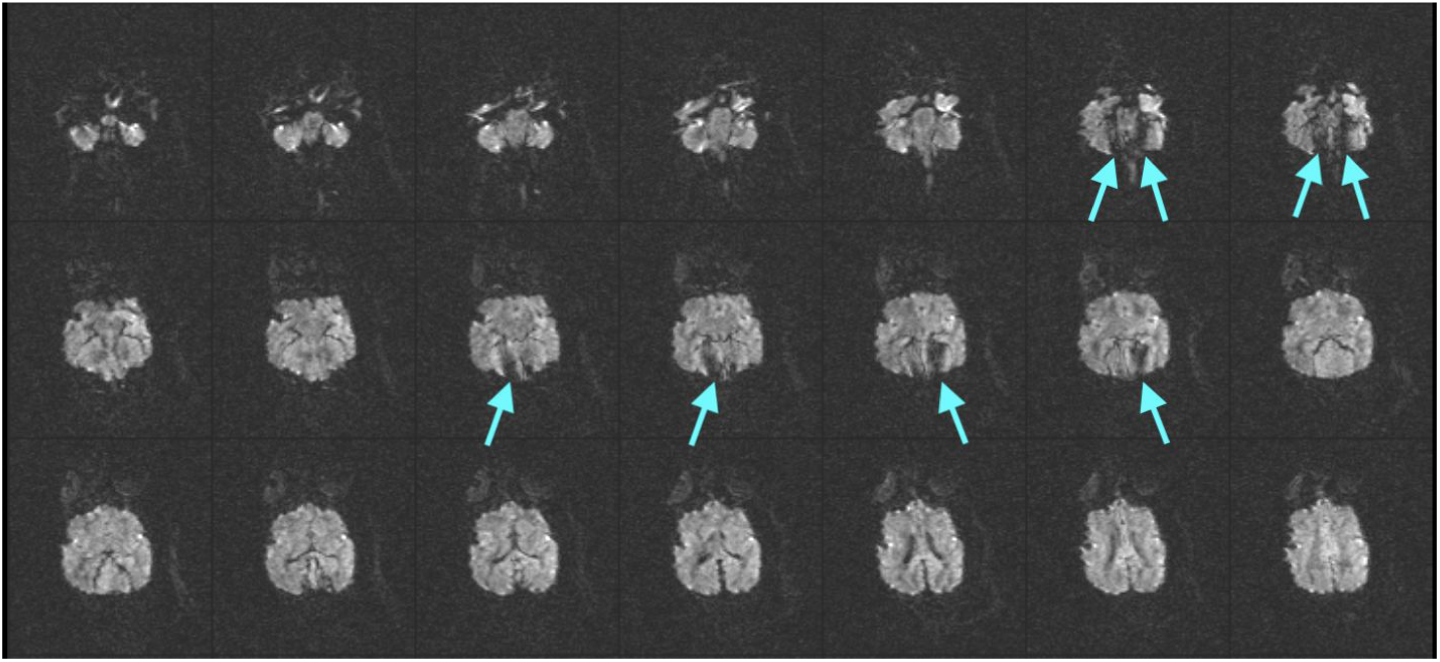
Exemplar diffusion-weighted images from yearling CSL Tarao, showing signal voids (arrows) that varied with slice position and dMRI direction in a random manner.

Subsequent review of the earlier data sets revealed that we had in fact encountered weaker, less frequent non-stationary signal voids in the DW images acquired from both Stern and Cronutt. The initial suspicion was of a mechanical resonance in the patient bed producing a DW direction-specific signal instability at b=1000 s/mm^2^. While this is a known problem on Siemens Trio scanners (Gallichan *et al*. 2010, Berl *et al*. 2015), it has rarely been an issue on the BIC scanner and later tests suggested that mechanical resonances were not the likely culprit.

Gallichan *et al*. (2010) found that signal voids can arise because of phase shifts in the partial Fourier sampling used in the phase encoding dimension. Mechanical resonances are only one possible cause. Without a good explanation for the origin of the voids, we nevertheless tried schemes to alleviate them. A simple tactic is to reverse the polarity of the phase encoding gradient, *e.g.* from L-R to R-L. We tested reversed phase encode gradient polarity on yearling CSL Hammond but aperiodic signal voids were still detected. We next tried b=850 s/mm^2^ at TE=90 ms and then b=1500 s/mm^2^ at TE=100 ms, maintaining 6/8ths partial Fourier for both L-R and R-L phase encode polarity. Aperiodic signal voids persisted. The voids were most common in cerebellum, suggesting that perhaps the endotracheal tube was causing a signal instability via magnetic susceptibility gradients. Test scans were next acquired on Hammond with the endotracheal tube removed and a nose mask to maintain anesthesia, but the non-stationary voids were still observed.

At this point, we realized that the most likely explanation was a shift of the resonance frequency caused by inflation of the animal’s chest with the intermittent manual ventilation. Modulation of the main magnetic field across the brain during respiration is a well-known source of physiological variance in human fMRI scans (Van de Moortele et al., 2002). In the case of our dMRI scans, the frequency shift due to movement of the sea lion’s chest may be sufficient to produce a phase shift in the partial k-space and cause the signal voids. This leaves two options: exchange partial Fourier for GRAPPA acceleration to maintain the TE below 100 ms and risk exchanging one artifact type for another, or change the procedure for monitoring and ventilating the animal during the dMRI scan. In future tests we will ventilate manually only during b=0 scans and we will include additional b=0 scans if extra time is required to complete the procedure. In spite of a few weak signal voids, we decided to process the 64-direction data set from CSL Stern and assess the tractography pipeline for basic structural connectivity estimation.

## Results

### Anatomical scans

Examples of high quality MP-RAGE, SPACE and T_2_-FLAIR images are shown in Figures 8, 9 and 10, respectively. Gray and white matter are clearly differentiated in the MP-RAGE and CSF appears dark, as in the human brain (Fig 8). The CSF is hyperintense in the SPACE image, with gray matter brighter than white matter (Fig 9). In the T_2_-FLAIR (Fig 10), normal tissue appears relatively isointense and CSF is dark. In some animals with suspected DOM toxicosis, hippocampal gliosis was detected easily in the T_2_-FLAIR, with corresponding atrophy evident in the MP-RAGE and SPACE scans (data not shown). An experienced ACVR board-certified veterinary radiologist (SD) determined that a full clinical read was feasible using the 3D scans.

**Figure 8.**
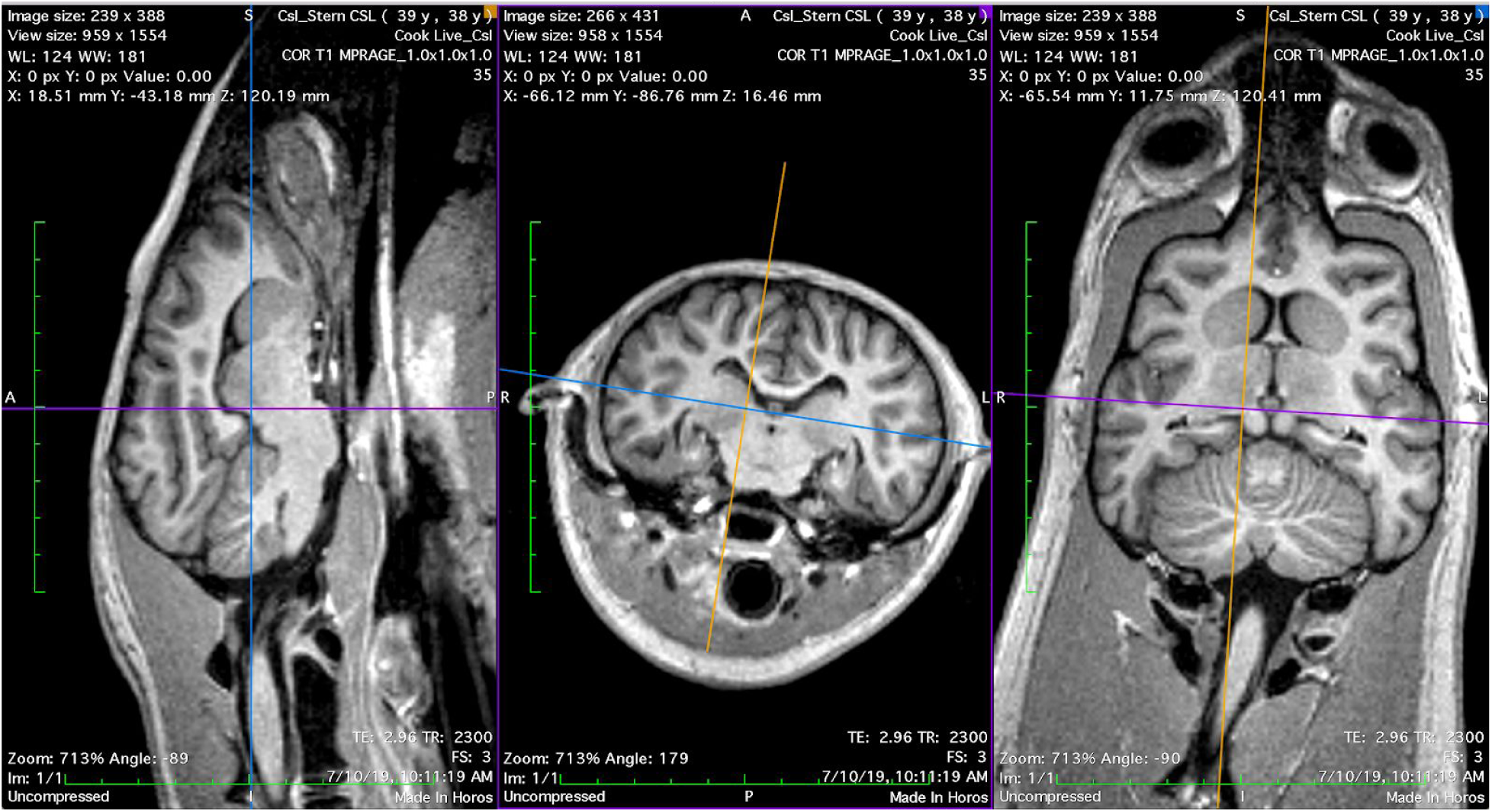
T_1_-weighted 3D MP-RAGE images obtained on yearling CSL Stern. Image orientation terminology: Left panel = sagittal, middle panel = transverse, right panel = dorsal.

**Figure 9.**
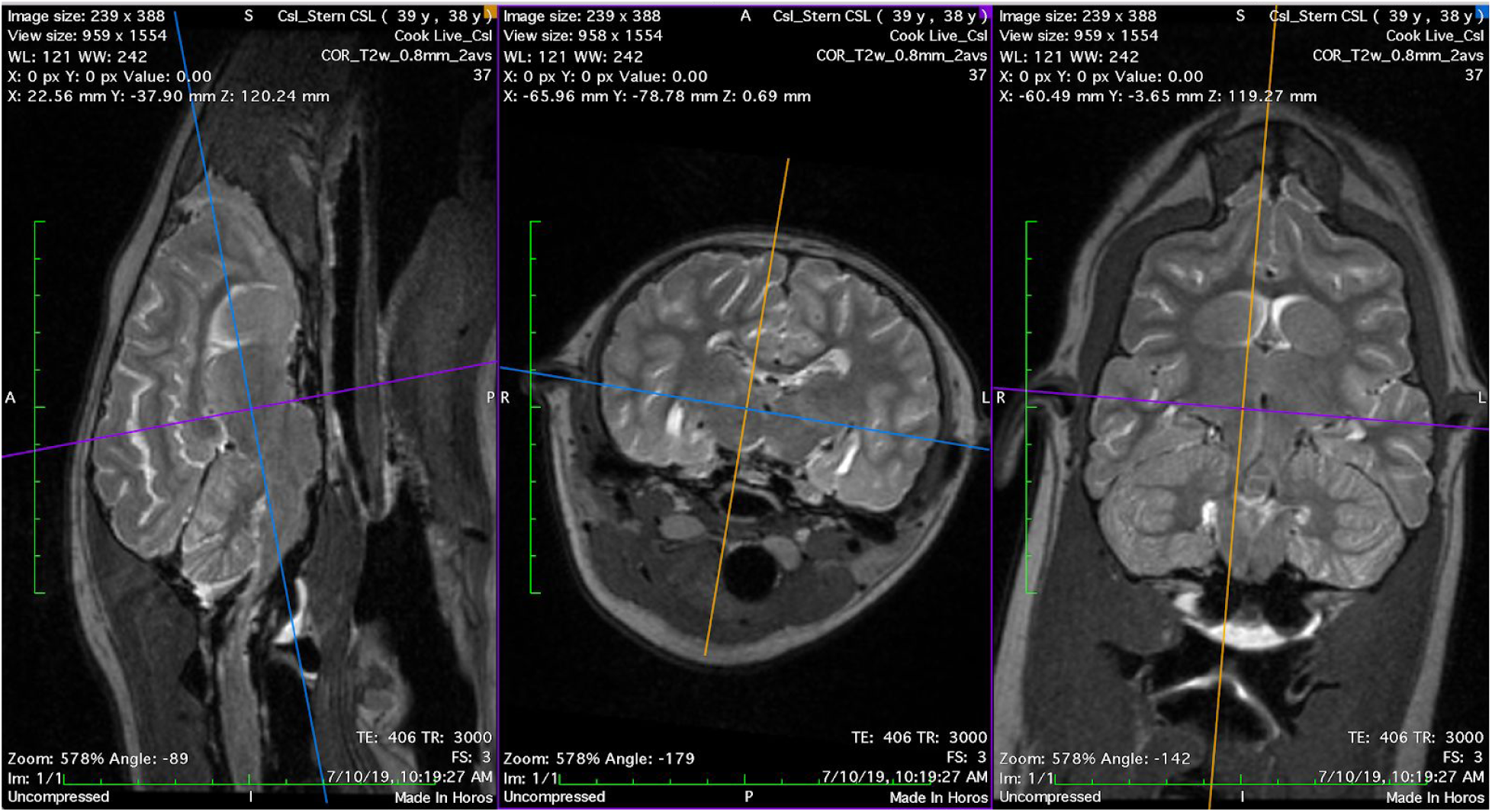
T_2_-weighted 3D SPACE images obtained on yearling CSL Stern.

**Figure 10.**
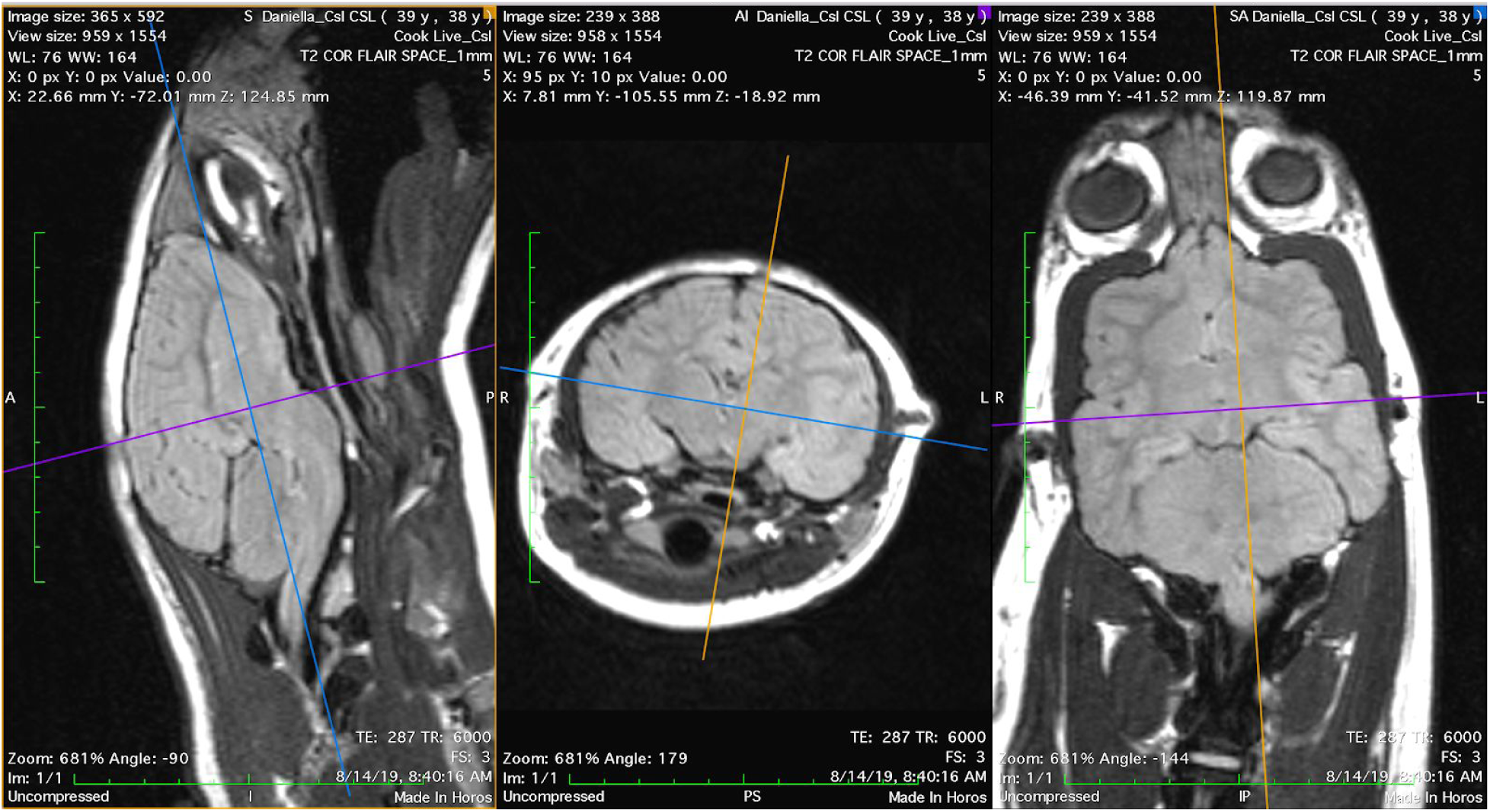
T_2_-weighted 3D FLAIR images obtained on yearling CSL Daniela.

Left and right hippocampi were segmented separately and volumetric values were obtained using the open-source software *3DSlicer* (www.slicer.org) for CSL Stern using T_2_ images (Figure 11). For segmenting hippocampal gray matter, an oblique orientation was selected perpendicular to the septo-temporal (longitudinal) axis of the hippocampus. This orientation has been used in prior hippocampal volumetry with sea lions (Montie *et al*. 2009, 2010, Cook *et al*. 2015), and allows clear contrast between the caudoventral boundaries of hippocampal gray matter, particularly in the *cornu Ammonis* (CA), and the CSF of the lateral ventricles. Tissue segmentation was cross-referenced on each oblique slice against a sagittal view. Hippocampal tracings included the CA subfields, dentate gyrus and alveus, as well as a portion of the subiculum. There are no clear anatomical criteria for distinguishing subiculum from CA in the sea lion. Septal and temporal boundaries of the hippocampus were based predominantly on the visible extension of hippocampal tissue in the sagittal view. Hippocampal tissue was clearly visible two to three slices temporal to the last oblique slice in which the cerebral peduncles could be viewed. The septal boundary was just dorsal to the dorsal boundary of the splenium of the corpus callosum. At this level in the oblique images the sea lion hippocampus appears flattened, as in primates. Sea lions do not have a large dorsal extension of CA as seen in dogs and rodents.

**Figure 11.**
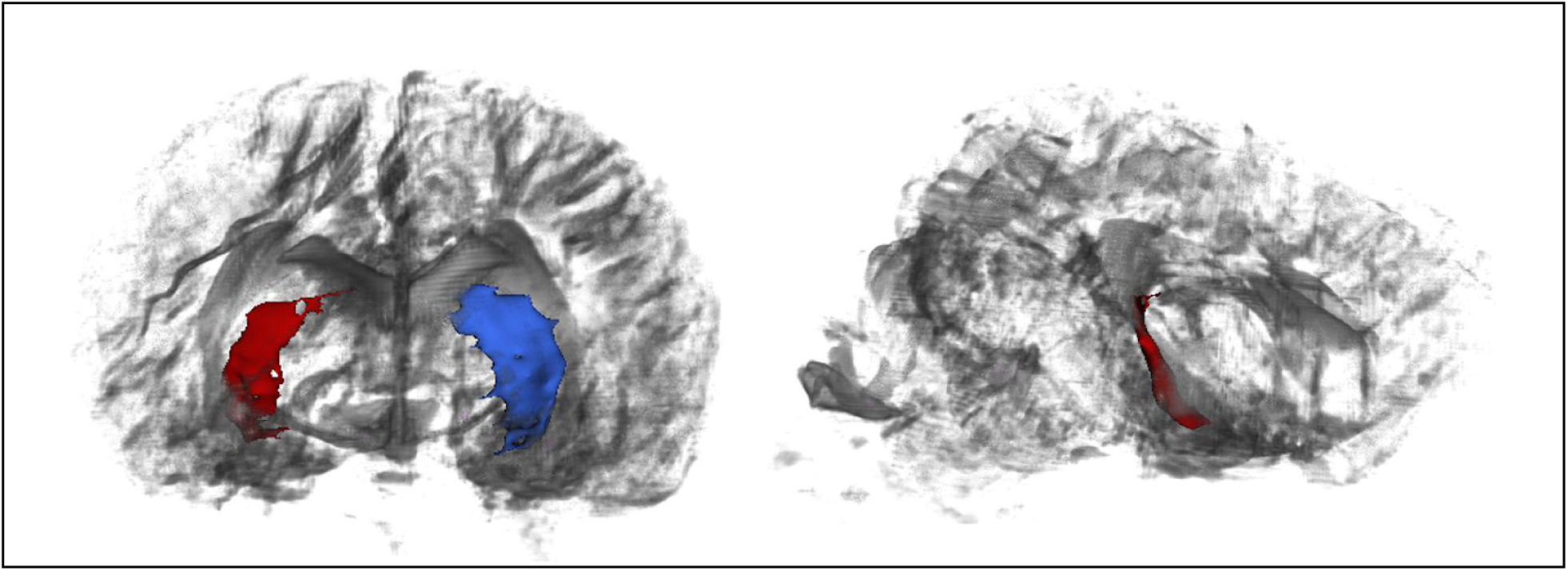
Left: Transverse render of right (red) and left (blue) hippocampal segmentations in a semi-transparent brain volume. Right: Sagittal render of right hippocampal segmentation. Both from CSL Stern.

The lateral boundary of the hippocampus was determined by the lateral ventricle of the temporal horn. Alveus was included where it could be clearly distinguished, but fimbria was not. The medial boundaries were set as the medial-most extension of subicular cortex. Rostroventral boundaries were defined by an arbitrary straight line connecting the medial-most portion of subiculum to the medial-most portion of parahippocampal tissue dorsal to parahippocampal gyrus.

Hippocampal volumes were computed using the *segmentstatistics* module in *3DSlicer*. Left hippocampus, as traced, was 580.2 mm^3^. Right hippocampus was 367.3 mm^3^. The notable difference in volumes matches radiological assessment of Stern’s images, which suggest enlarged ventricles around the CA of the right hippocampus. Hippocampal volumes measured here are on the lower end of the range of previous volumetry in sea lions. The differences may be due to the young age of the animal and potential bilateral atrophy. In addition, unlike previous segmentations with CSLs, high quality 3D imaging allowed us to carefully trace in an oblique plane while also using sagittal orientation to carefully identify the longitudinal boundaries of hippocampal tissue. Montie *et al*. (2010) previously measured hippocampal volumes in six sea lions in the ∼700-1000 cm^3^ range using 2D T_2_-weighted imaging with 0.3 mm x 0.3 mm resolution in-plane and 2.5 mm thick slices. Follow-up work by Montie *et al*. (2012) using the same 2D acquisition identified similar volumes in fifteen further sea lions. Cook *et al*. (2015) measured right and left hippocampus in 30 sea lions using 2D T_2_-weighted imaging with 0.695 mm x 0..695 mm resolution in-plane and 2.0 mm thick slices, and found volumes ranging from ∼400 to 900 cm^3^. Some of these animals had severe hippocampal atrophy. General segmentation criteria were similar in all cases. In the current work, with isotropic 3D imaging and elimination of CSF signal using FLAIR, we were able to delineate all likely hippocampal tissue along the septotemporal axis but medial hippocampal tissue boundaries were not always clear. Cook *et al*. (2018) measured hippocampal volumes from high-resolution postmortem MRI (0.6 mm x 0.6 mm x 0.5 mm) and found higher volumes, ranging from ∼900 to 1800 cm^3^. Given the limited prior work describing sea lion hippocampal anatomy, the present imaging protocols provide an opportunity to carefully delineate and standardize measures using anatomical criteria.

### Functional MRI

Functional connectivity analysis was conducted on data obtained from CSL Tarao. Data preprocessing was carried out using a combination of manual steps and FEAT (FMRI Expert Analysis Tool) Version 6.00, part of FSL (FMRIB’s Software Library, www.fmrib.ox.ac.uk/fsl) (Woolrich et al. 2001). The following corrections were applied in this order: (i) volume realignment for motion correction using MCFLIRT (Jenkinson et al. 2002); (ii) slice-timing correction using Fourier-space time-series phase-shifting; (iii) non-brain removal by multiplication with a hand-drawn brain mask; (iv) spatial smoothing using a 2mm Gaussian kernel; (v) grand mean intensity normalisation of the entire 4D dataset by a single multiplicative factor; (vi) high-pass temporal filtering (Gaussian-weighted least-squares straight line fitting, with sigma=75.0s).

Next, we sought to assess systematic effects of the manual ventilation procedure using an independent component analysis (ICA)-based exploration of the time series with automatic dimensionality estimation using MELODIC (Beckmann & Smith 2004). Independent components were manually evaluated with the grading tool *PICAchooser* (https://github.com/bbfrederick/picachooser) (Frederick 2020) to flag artifactual components at the manual ventilation rate. Fourteen out of 21 independent components found by MELODIC exhibited well defined, pseudo-periodic maxima across the time series every 50-60 seconds. The maxima had differing shapes and temporal lags for each IC but maintained the same frequency of ∼0.02 Hz. The spatial locations of the ICs were most prominent in inferior brain regions and locations consistent with large arteries, suggesting a vascular origin for the signal changes (Figure 12). Non-stationary blood gas concentrations, in particular arterial CO_2_, modulate arterial tone in a manner that produces concomitant BOLD signal changes (Wise *et al*. 2004). The fourteen components assigned as artifacts of the manual ventilation were removed from the preprocessed data using “fsl_regfilt” prior to further analysis. In total, these components accounted for 38.4% of total signal variance.

**Figure 12.**
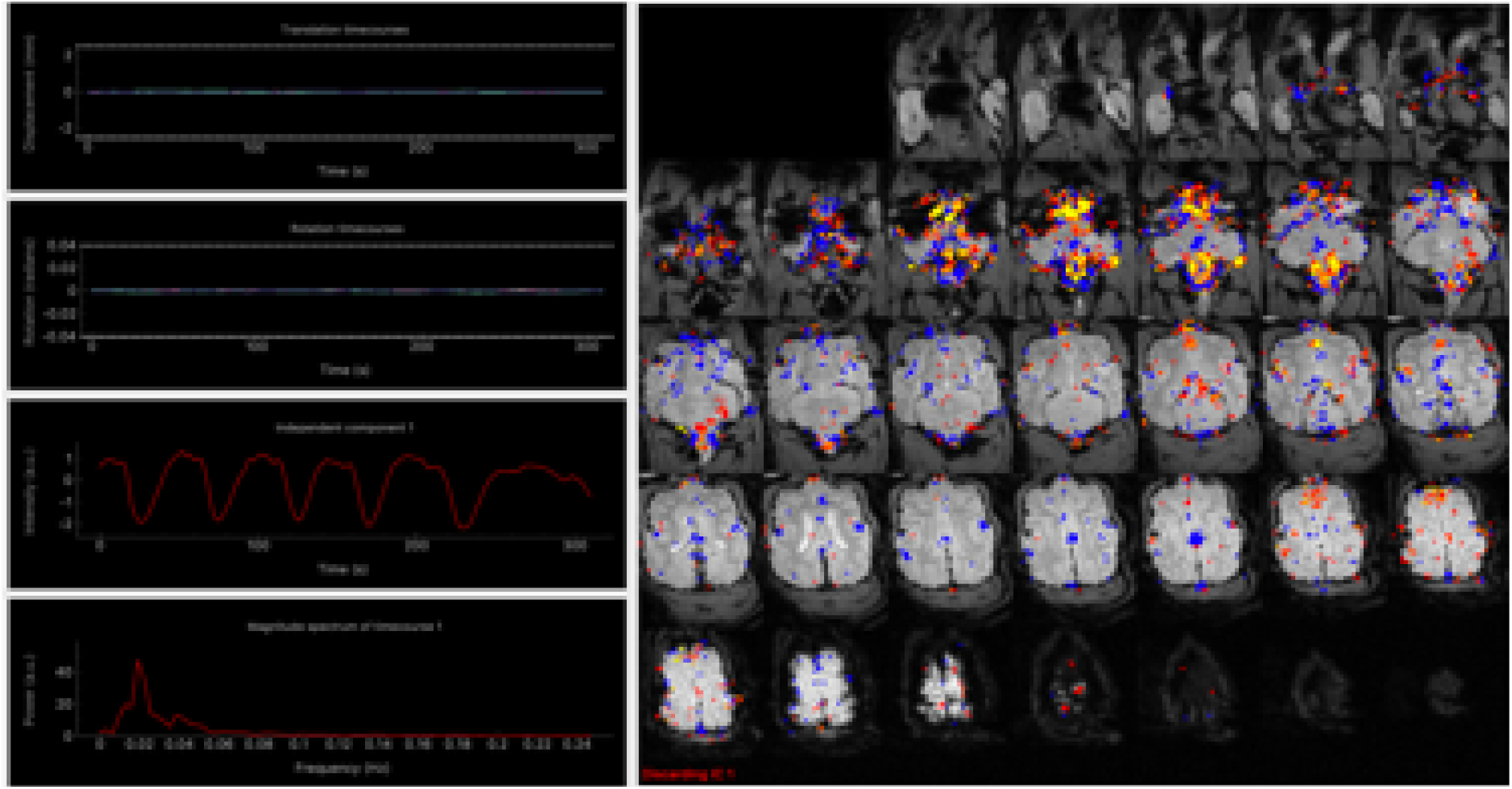
The first of 14 independent components identified using MELODIC independent component analysis, sorted using *PICAchooser*. The third row of the left-hand column shows the time series corresponding to the colored voxels displayed in the right panel. The six maxima of the time series correspond to the manual ventilation, approximately one per minute. The brightest voxels arise at the base of the brain and brainstem and around the known locations of large vessels, especially arteries.

For functional connectivity analysis of the cleaned data, a dorsal left hippocampal mask was created on the T_1_-weighted structural image in FSLEyes, following Cook *et al*. (2015). White matter and CSF masks were also generated from the structural image using FSL’s FAST (Zhang *et al*. 2001), and thresholded at p(tissue) > 0.95. Hippocampal, white matter, and CSF masks were registered into native functional space using FSL’s FLIRT (Jenkinson & Smith 2001, Jenkinson *et al*. 2002). A time course from the pre-processed data was extracted from each registered mask, computed as the average BOLD signal at each time point for each region.

Temporal correlation analysis was conducted in FSL’s FEAT using the general linear model with the dorsal left hippocampal time course as the primary covariate. The white matter and CSF time courses (Behzadi *et al*. 2007) plus the six realignment parameters (three translations, three rotations) were included as nuisance covariates to mitigate physiological noise and motion. Resulting correlation maps were registered to anatomical space for display, to indicate areas with highest functional connectivity with the dorsal left hippocampus (Figure 13).

**Figure 13.**
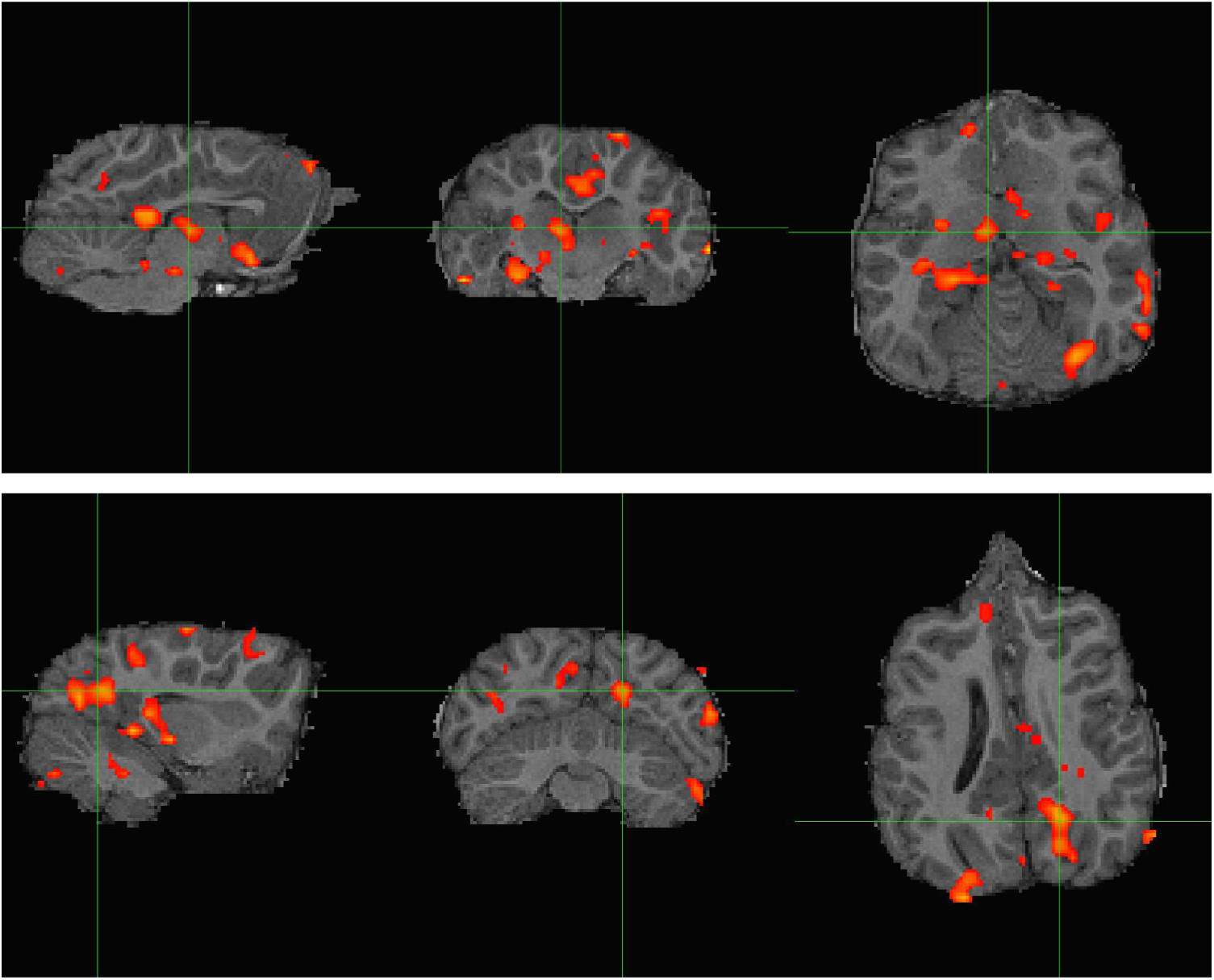
Resting-state connectivity analysis for CSL Tarao presented in, from left to right, sagittal, transverse, and dorsal planes. Top: Z-score map thresholded at Z > 3.1 computed from a dorsal left hippocampal seed. Strongest correlation was observed in the cingulate cortex and dorsal thalamus (cross hairs), ventral caudate nuclei, and right hippocampus (contralateral to the seed). Bottom: Another view of connectivity with the dorsal left hippocampal seed, presenting an area of caudal cortical activation likely overlapping with visual processing regions (cross hairs).

As in Cook *et al*. (2015), there was significant co-activation with the dorsal hippocampus in dorsal thalamus and the caudate nuclei (Figure 13, top). This is in line with established evidence regarding hippocampal connectivity in humans and other mammals (Stein *et al*., 2000). Work in human epilepsy indicates that hippocampal-thalamic pathways can be affected by chronic seizures (Barron *et al*., 2014). Contralateral hippocampal coactivation was also observed. Unlike in Cook *et al*. (2015), there was significant correlation in posterior dorsal cortex, potentially in a visual processing region (Figure 13, bottom), which is in line with evidence on hippocampal connectivity in rodents and humans (Eichenbaum, 2000; Lavenex & Amaral, 2000, Ranganath *et al*. 2005). However, the presence of high coactivation in the brainstem and parts of the lateral ventricles suggests the data are affected by residual physiological noise. Manual ventilation produces large magnetic field variations over the brain from the magnetic susceptibility of the abdomen, as well as the possibility of large direct motion of the head, and the denoising tactics employed here were likely insufficient. Future analyses might include covariates derived from direct physiological monitoring, in particular the motion of the chest which can be recorded with a plethysmograph belt, and principled removal of non-stationary arterial CO_2_ effects (Tong *et al*. 2019).

### Cerebral blood flow mapping

The ASL data from CSL Cronutt was analyzed following established methods for determining CBF in the human brain. The label-control time series was first realigned to mitigate effects of motion. Regional CBF was then computed using single compartment kinetics (Alsop *et al*. 2015). The model used an assumed brain/blood partition coefficient of 0.9 and labeling efficiency of 0.72 (Dolui *et al*. 2019), determined based on experiments conducted in normal humans. The T_1,blood_ = 1.74 s was calculated from the relationship 1/T_1_= 0.62.Hct + 0.28 (Li *et al*., 2017) using a hematocrit value of 0.47 obtained from blood drawn under anesthesia during the scanning session.

A whole brain mask was determined from Cronutt’s 3D T_2_-weighted image using FSL’s *BET*. The masked CBF map is shown overlaid on the T_2_-weighted anatomical scan in Figure 14. The gross anatomical features in the CBF map - gray matter CBF approximately three times greater than white matter - are recognizable features seen in the normal human brain. In spite of possible species differences and the effects of anesthesia, it is instructive to compare the global mean CBF found for Cronutt with typical human data obtained using ASL. The mean global CBF in Cronutt was 81.3 ml/100 g/min. This compares favorably with values of around 80 ml/100 g/min measured in 8 year-old humans (Satterthwaite *et al*. 2014), using as a crude index the age of sexual maturity, with male sea lions reaching breeding age at around 5-6 years old.

**Figure 14.**
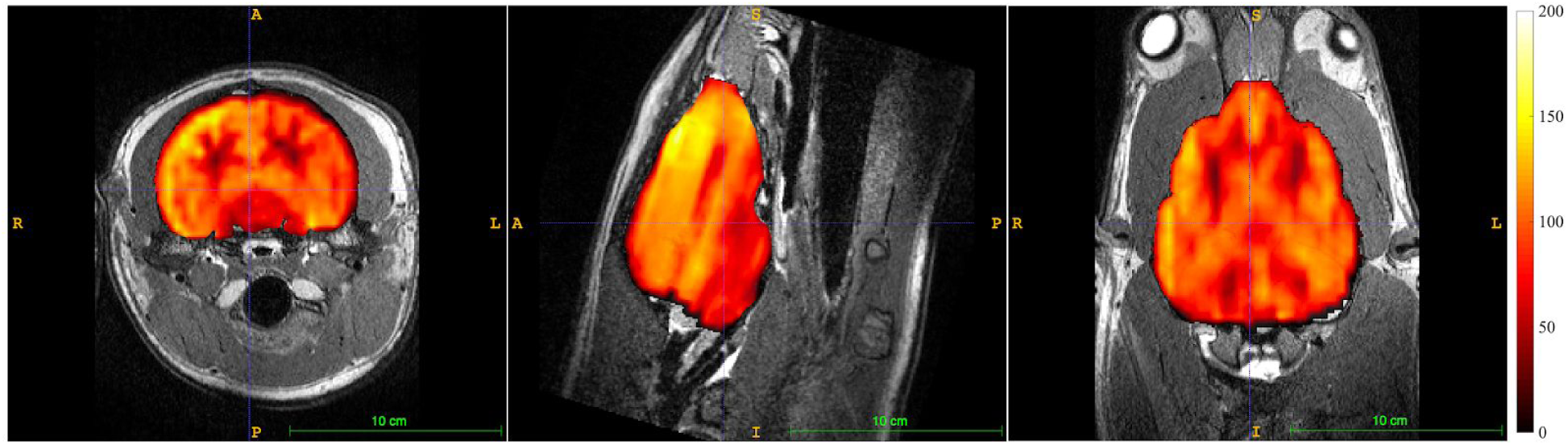
Cerebral blood flow map obtained from 3 year-old CSL Cronutt using a 2-shot stack-of-spirals PCASL sequence with 1800 ms labeling period and 1200 ms post-label delay. Color scale units are ml blood/100 g tissue/minute. The CBF map is overlaid on the T2-weighted anatomical scan. Images are, left to right, in the transverse, sagittal, and dorsal planes.

### Diffusion imaging

The dMRI data obtained on CSL Stern had the fewest non-stationary signal voids that could be determined by inspection, and these images were submitted for tractographic analysis. We noted only four possible signal voids, all confined to the cerebellum, in the entire dMRI data set for Stern.

Processing of dMRI data was performed using FDT (FMRIB’s Diffusion Toolbox, Oxford University, UK). Images were first converted from DICOM to NIFTI format using *DCM2Nii*. The six b=0 images were co-registered and averaged to serve as a structural reference for the dMRI data. The non-canonical orientation of the sea lion’s head during imaging required a correction to the DW vectors produced by the scanner: the y and z axes were swapped, then the signs were reversed for the x and y axes. The results of the corrections were checked carefully in all three planes using FA maps with voxelwise principal diffusion directions overlaid, all generated via FSL’s *DTIFit* tool. Diffusion data were then registered to the average b=0 image. A brain mask generated by FSL’s *BET* tool, with manual editing, was created from the average b=0 image and used to skull-strip the registered diffusion data. Simultaneous eddy current correction and volume realignment for motion correction was then performed on all DW images using FSL’s *eddy* tool. Corrected images were used henceforth.

Images of FA and MD were produced using FSL’s *DTIfit* tool. This utility assumes a tensor model for the underlying diffusion (Basser *et al*., 1994). For tractography, however, we used a probabilistic model to estimate local diffusion parameters in each voxel with FSL’s *bedpostx*, a Bayesian estimation probabilistic diffusion model which attempts to account for crossing fibers by Markov Chain Monte Carlo sampling to build up distributions on diffusion parameters within each voxel.

The output from *bedpostx* (Figure 15, top) was used to perform probabilistic fiber tracking using FSL’s *probtrackx* (Behrens *et al*., 2003, 2007). As in Cook *et al*. (2016), we traced the crus and pillar of fornix tracts. Hand-drawn anatomical seeds for these tracings were created in FSL’s *FSLView* directly in DTI space, over FA maps. The seed was placed in the crus of the fornix and situated in high FA apparent white matter with clear rostro-caudal diffusion directionality per an overlaid diffusion vector map. The seed was just ventral to the corpus callosum, and the rostral boundary was caudal to the ventral turn of the rostral fornix. Termination masks were also created, one in the dorsal plane ventral to the splenium and one in the transverse plane caudal to the septum, which terminated any apparent tracts entering the corpus callosum and septum pellucidum, respectively. Tracing was conducted with 5,000 streamlines per seed voxel, a step length of 0.5 mm, and a standard curvature threshold of 80 degrees. The resulting tracts thresholded at 1% clearly demarcated the fornix.

**Figure 15.**
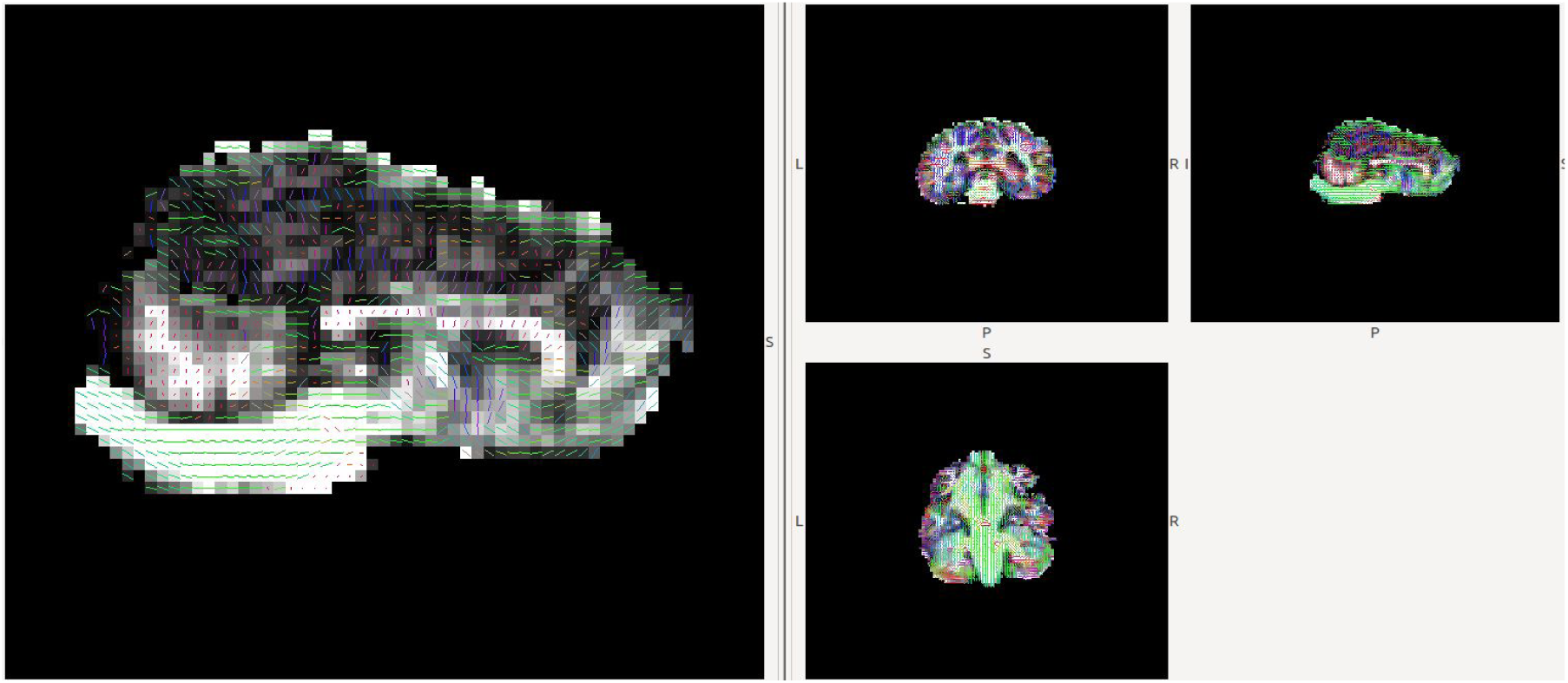
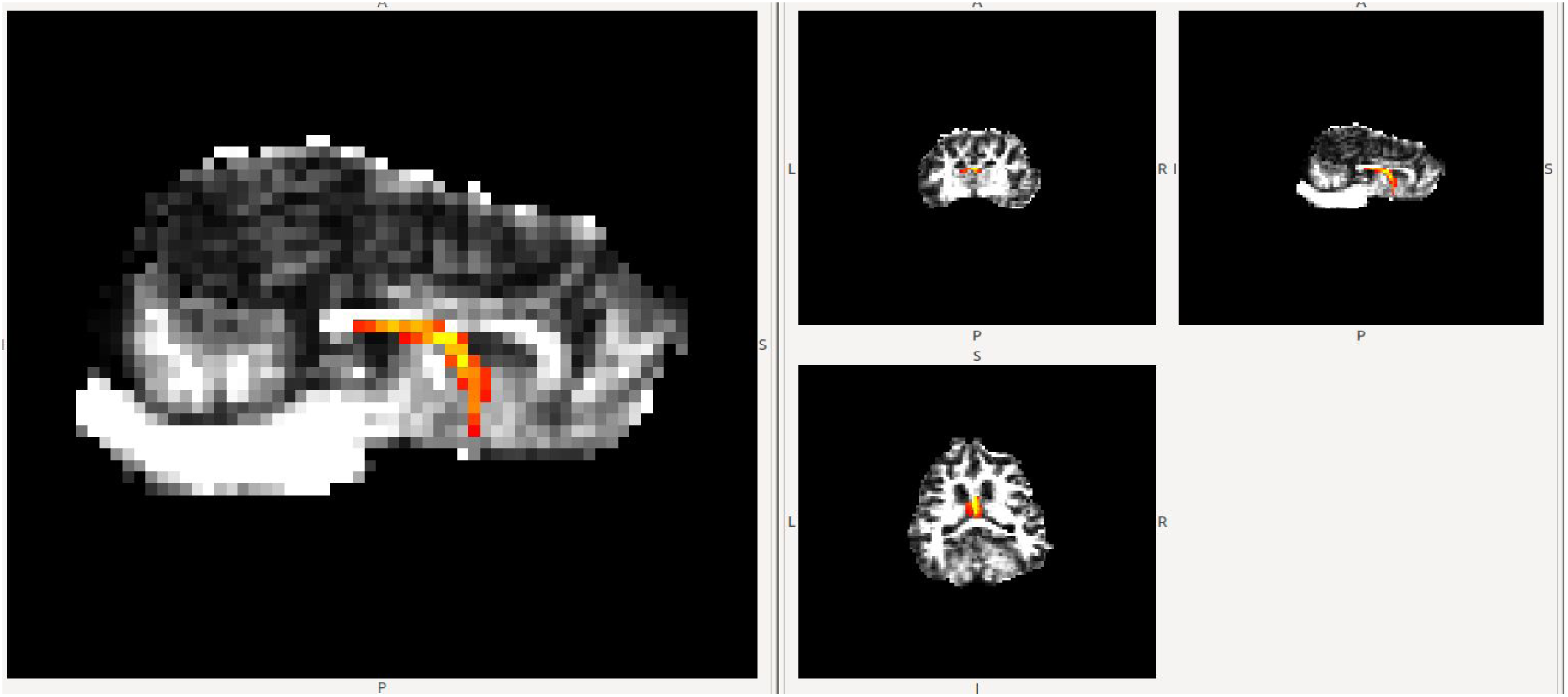
Top: Voxel-by-voxel representation of principle diffusion directions in sagittal (left, large) and all three canonical (three right, small) planes, shown in diffusion space/resolution. Diffusion data are overlaid on a b0 image. Each voxel has a line and color indicating principle water diffusion direction. Green voxels are rostro-caudal, red voxels left/right, and blue voxels dorso-ventral. Bottom: Hippocampal fornix traced from a bilateral fornix pillar seed, thresholded at 1% of total streamlines.

Both probabilistic and deterministic fornix tracings in the current study were in line with previous post mortem DTI analyses in sea lions (Cook *et al*. 2018), suggesting that Stern’s dMRI data were acceptable despite the aperiodic signal voids noted in cerebellum. Fornix was clearly identified in FA maps based on anatomical criteria with principle diffusion maps benefitting seed placement (Figure 15, bottom). The apparent lack of traceable forniceal white matter in the left hippocampus (Figure 16) may be indicative of damaged tissues. In Cook *et al*. (2018), fornix FA values were significantly lower in animals with diagnoses of DOM toxicosis and hippocampal atrophy, as also reported in human patients with mTLE (Concha *et al*., 2010). Deterministic and probabilistic tracing rely in part on FA thresholds voxel-by-voxel. However, partial voluming around ventricles has been identified as a confound in DTI of the fornix previously (Kuroki *et al*., 2006). Whether the present results are representative of white matter pathology will require resolution of the signal void issue and, ideally, a comparison of *in vivo* to post mortem dMRI from the same brain to confirm.

**Figure 16.**
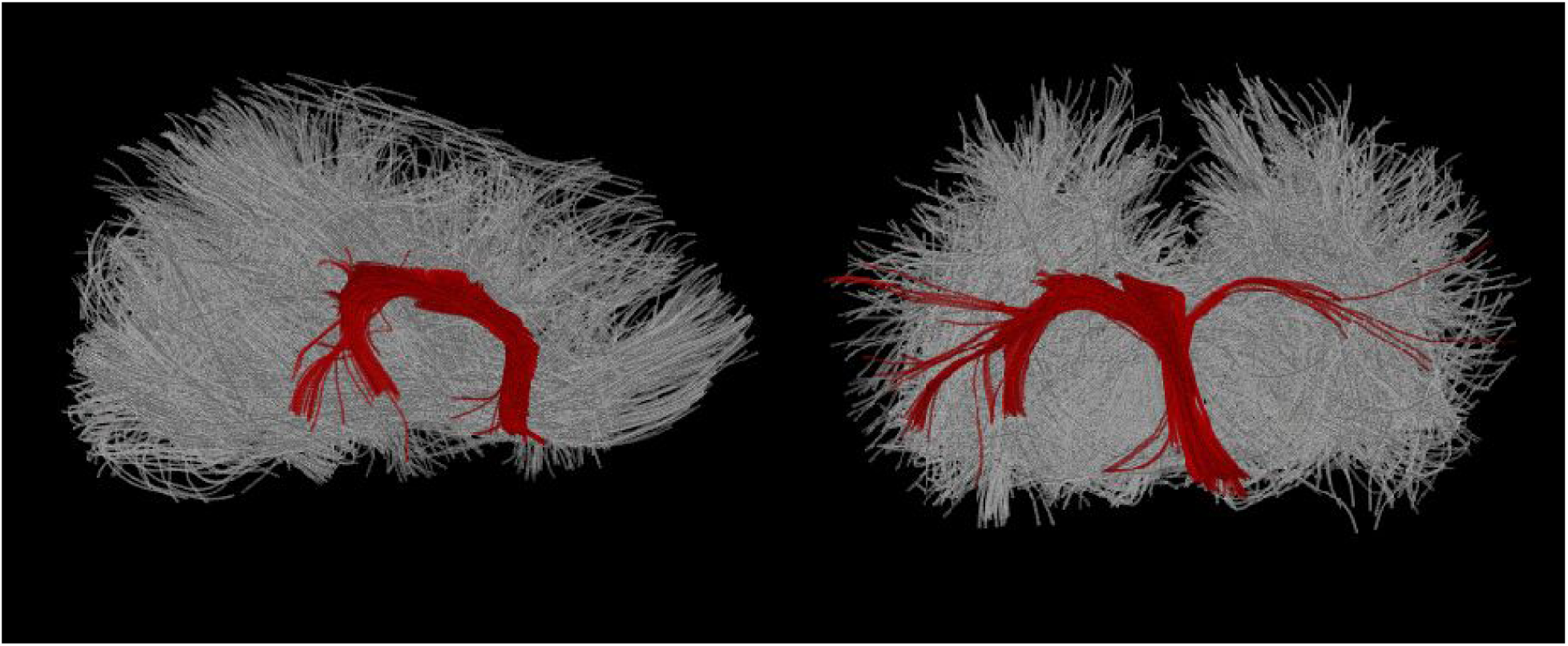
A deterministic tracing of hippocampal fornix produced from a fornix pillar seed analogous to that used in Figure 15, overlaid on a whole brain rendering of all fibers, in the sagittal (left) and transverse (right) views. Deterministic tracing is not thresholded but termination masks were used dorsal to the dorsal fornix and caudal to the caudal-most portion of fornix.

## Discussion

We have developed a research MRI protocol suitable for studying California sea lions suspected of domoic acid toxicosis. A list of the main scan parameters appears in Table 2. The scan protocol is suitable for pinnipeds over a range of sizes, having been developed on animals of 21-96 kg. Our long-term goal is to study the development of sea lions exposed *in utero* to domoic acid, from 6 months to 4 years of age. Beyond four years it is likely that male animals would be too large for our MRI scanner. The availability of a wide-bore scanner with a patient bed rated to 250 kg would permit scans of slightly larger animals, assuming suitable modifications to the procedure for transfering to and from the MRI bed.

**Table 2:** Main MRI scan parameters.

| Scan type | Sequence type | TR | TE | Spatial resolution (mm) | Additional parameters | Acquisition time | Notes |
| --- | --- | --- | --- | --- | --- | --- | --- |
| T1-weighted anatomical | 3D MP-RAGE | 2300 ms | 2.96 ms | 1.0 x 1.0 x 1.0 | T<sub>INV</sub>= 900 ms, non-selective inversion. Excitation flip angle = 9 degrees. | 7m 23s | *a* |
| T2-weighted anatomical | 3D SPACE | 3000 ms | 406 ms | 0.8 x 0.8 x 0.8 | Turbo factor = 141, slice turbo factor = 2. | 6m 02s | *b* |
| T2-weighted anatomical with fluid attenuation | 3D FLAIR-SPACE | 6000 ms | 287 ms | 1.0 x 1.0 x 1.0 | T<sub>INV</sub>= 2200 ms, non-selective inversion. Turbo factor = 101, echo trains per slice = 2. | 8m 26s | *c* |
| BOLD-weighted functional MRI | 2D GRE-EPI | 2000 ms | 25 ms | 2.5 x 2.5 x 2.5 | No. of volumes = 160. Excitation flip angle = 65 deg. Slice order = descending. Read bandwidth = 2192 Hz/pixel, echo spacing = 0.52s. | 5m 22s | *d* |
| CBF mapping | 3D stack-of-spirals PCASL | 4600 ms | 8.7 ms | 3.0 x 3.0 x 3.0 (or 3.8 x 3.8 x 3.8) | Labeling duration = 1800 ms, post-labeling delay = 1200-1800 ms | 4m 37s | *e* |
| Diffusion MRI | DW-SE-EPI | 5800 ms | 93 ms | 2.0 x 2.0 x 2.0 | 64 directions at b = 1000 s/mm<sup>2</sup> plus six b=0 images | 6m 58s | *f* |

*a* Excellent gray/white matter contrast; useful for quantitative morphometry.
*b* Good tissue contrast & visualization of CSF. Also shows edema & gliosis.
*c* Excellent identification of pathology, especially edema & gliosis. Can be used for quantitative morphometry.
*d* FA < 65 deg would reduce physiological artifacts without compromising intended BOLD contrast.
*e* For 3.0 mm or 3.8 mm voxels a 4- or 2-shot segmented spiral readout is used, respectively. Optimal PLD may vary depending on animal size/age.
*f* Requires 6/8ths partial Fourier to keep TE < 100 ms, which may enhance sensitivity to respiratory artifacts.

Clinical evaluation of DOM toxicosis has used 2D MRI scans previously. We opted to use 3D anatomical scans because these permit improved tissue volume measurements, as required for a quantitative longitudinal study of changes following DOM toxicosis, and for arbitrary re-slicing post hoc. An experienced ACVR-board certified veterinary radiologist (SD) confirmed the utility of the 3D scans for clinical purposes. Beyond DOM toxicosis, we predict that the 3D scans would permit clinical discrimination of brain pathologies such as tumors, hemorhages and the presence of gas bubbles.

The greatest technical challenges were caused by respiration, in particular the requirement to manually ventilate the animal periodically to avoid hypercapnia. Respiration negatively affected all scan types when using standard human parameters, necessitating scan-specific tactics to ameliorate. In 3D anatomical scans, respiration presented a challenge due to the orthogonal orientation of the sea lion’s brain relative to a supine human, and the comparatively large volume of tissue in a sea lion’s throat; normal respiration produces very small movements of throat signal in humans. A comparable problem in human imaging is swallowing, which produces large excursions in both A-P and H-F directions and would cause analogous errors were it to occur with the frequency of respiration.

Our test of functional connectivity from fMRI yielded results at 3 T that matched prior results obtained at 1.5 T on sea lions, with dorsal thalamus and dorsal caudate co-activated with a dorsal hippocampus seed. In general, the magnitude of BOLD contrast is expected to increase with B_0_, offering more sensitivity at 3 T, but this advantage manifests only if physiological changes unconnected to neurovascular coupling can be minimized (Liu 2016, Wald & Polimeni 2017). We observed a major perturbation consistent with manual ventilation. Many regions, notably arteries in the inferior slices, fluctuated at the pseudo-periodic rate of the manual ventilation and exhibited high signal variance, strongly suggesting a BOLD signal change due either to non-stationary arterial CO_2_ (Tong & Frederick 2010, Tong *et al*. 2019) and/or non-stationary deoxyhemoglobin concentration in arterial blood whenever SaO_2_ departs from the assumed value of 100% (Aso et al. 2019). Such respiration-induced changes are typical in humans. The issue here is the magnitude of the response to intermittent manual ventilation. In future, during fMRI acquisition we will attempt a more regular manual or mechanical ventilation with smaller volumes, to better resemble normal ventilation. A scripted ventilation routine will also give us ground truth knowledge to improve identification and separation of respiratory effects from ongoing brain activity.

We do not yet know whether manual ventilation affected the CBF map measured with ASL, via the well-known vasodilative effects of arterial CO_2_. In future tests we will compare CBF maps produced with large volume, low rate ventilation as used here to a scripted procedure of shallower, more frequent ventilation as will be used during the fMRI acquisition. We will also conduct more tests of labeling duration and post-labeling delay. Older or larger sea lions may benefit from a longer PLD if they have greater arterial transit times. However, longer arterial transit could be offset by the higher hematocrit (*i.e.* shorter blood T_1_) expected in healthy older animals. Furthermore, the assumed brain/blood partition coefficient and blood T_1_ estimates derive from human work that may not be appropriate for sea lions. The range of hematocrit values we observed across seven animals was large (0.28-0.47; median 0.40), in part due to age (7 months to 3 years) and overall health, but also perhaps because of differing splenic responses to anesthesia. In future, a multiple PLD approach may prove necessary for studying developing sea lions because the ATT may change with the animal’s size, while the CBF itself may change with age. Sea lions in the range 6 months to 4 years correspond approximately to children from 1 to 14 years old. The CBF has been observed to be high in human infants, around 100 ml/100 g/min, dropping to ∼80 ml/100 g/min by 8-9 years old (Suzuki *et al*. 1990, Satterthwaite *et al*. 2014). By age 18 the global CBF drops further, to a typical adult value of ∼60 ml/100 g/min. In adulthood there is then a steady decrease to around 40 ml/100 g/min by age 70 (Dolui *et al*. 2016, Dolui *et al*. 2017). Whether the sea lion brain follows a similar trajectory is an area for future research.

The manual ventilation appears to have caused a large artifact in our dMRI paradigm. Most likely the problem is due to chest motion - the through-space modulation of the magnetic field over the brain via chest magnetic susceptibility, and possibly some direct motion coupled to the head - rather than non-stationary blood gases. Presently, we acquire one non-DW image for every ten DW images as templates for eddy current and distortion corrections. In future, motion sensitivity can be reduced by ventilating during the non-DW images only, adding additional non-DW images to accommodate the ventilation if necessary, or perhaps with a reduced volume, higher rate ventilation procedure.

### Future developments

Improving the protocol beyond the current version would be feasible with a more sensitive receiver coil than the four-channel human neck coil. A ring or a helmet coil with eight or more channels would permit the use of GRAPPA acceleration in 3D anatomical scanning, perhaps allowing higher spatial resolution in an acceptable scan time, and would also permit the use of SMS acquisitions for both the fMRI and dMRI scans. It is feasible to use low SMS factors with the human neck coil, but our pilot tests indicated some artifacts which were not worth the modest speed gains. A custom coil, designed with the constraints of anesthesia and animal monitoring in mind, would be preferable.

We have not yet had the opportunity to test a T_2_*-weighted anatomical scan for susceptibility-weighted imaging (SWI) or quantitative susceptibility mapping (QSM). This sort of contrast is primarily used to detect hemorrhage but may also be useful for differentiating calcification, iron deposits and microbleeds (Liu *et al*. 2017). We would deploy the sequence in the event we observed pathology on one of the conventional anatomical scans. A future protocol might also include MP2RAGE for tissue and CSF segmentation (Wang *et al*. 2018), for enhanced deep gray matter contrast (Tanner *et al*. 2012, Okubo *et al*. 2016), or for detecting focal epileptogenic lesions (Kotikalapudi *et al*. 2019). The MP2RAGE could be a replacement for the MPRAGE we employed here. We will continue to test additional sequences as opportunities arise.

## Conclusions

The protocol presented in this paper establishes a framework for studying the neurological effects of exposure to toxic algae in developing sea lions. We propose that our protocol may also be a good starting point for MRI scanning of other marine mammals, including phocid seals, sea otters and perhaps even smaller cetaceans (dolphins and porpoises). While there is a large range of head and brain sizes and brain anatomies across marine mammals that might fit an MRI scanner, the tissue contrast is reasonably consistent across mammalian brains. Standard contrast parameters for human brain imaging were used here and were found to deliver high quality images. Respiration was the greatest impediment to scan quality, and each scan type required careful tuning to overcome the largest artifacts. Some further work is needed to mitigate the remaining consequences of manual ventilation in the fMRI and dMRI scans, and more experience is needed with CBF mapping to be able to fully assess data quality.

## Acknowledgements

We are indebted to the staff of The Marine Mammal Center, UC Berkeley veterinarians Greg Lawson and Christie Ferrecchia, and the staff of Six Flags Discovery Kingdom for their animal welfare expertise. We thank the Wheeler Family Foundation for their generous support of the Henry H. Wheeler, Jr. Brain Imaging Center, and the National Science Foundation for support through their Major Research Instrumentation Program, award number BCS-0821855.

